# Gambling environment exposure increases temporal discounting but improves model-based control in regular slot-machine gamblers

**DOI:** 10.1101/2021.07.15.452520

**Authors:** Ben Wagner, David Mathar, Jan Peters

## Abstract

Gambling disorder is a behavioral addiction that negatively impacts personal finances, work, relationships and mental health. In this pre-registered study (https://osf.io/5ptz9/?view_only=62956a2afcd1495984db4be18c54b50a) we investigated the impact of real-life gambling environments on two computational markers of addiction, temporal discounting and model-based reinforcement learning. Gambling disorder is associated with increased temporal discounting and reduced model-based learning. Regular gamblers (n = 30, DSM-5 score range 3-9) performed both tasks in a neutral (café) and a gambling-related environment (slot-machine venue) in counterbalanced order. Data were modeled using drift diffusion models for temporal discounting and reinforcement learning via hierarchical Bayesian estimation. Replicating previous findings, gamblers discounted rewards more steeply in the gambling-related context. This effect was positively correlated with gambling related cognitive distortions (pre-registered analysis). In contrast to our pre-registered hypothesis, model-based reinforcement learning was improved in the gambling context. Here we show that temporal discounting and model-based reinforcement learning are modulated in opposite ways by real-life gambling cue exposure. Results challenge aspects of habit theories of addiction, and reveal that laboratory-based computational markers of psychopathology are under substantial contextual control.

## Introduction

Gambling disorder is a behavioral addiction that can have detrimental effects on quality of life including personal finances, work, relationships and overall mental health (Blaszczynski & Nower, 2002; Muggleton et al., 2021). Despite these negative consequences, many gamblers are motivated to continue to play, and praise the temporary excitement and pleasure (Fauth-Bühler et al., 2017). Accumulating evidence suggests similarities of gambling disorder and substance-use-disorders both on behavioral, cognitive and neural levels (Balodis & Potenza, 2020; Leeman & Potenza, 2012; Lobo et al., 2015; N. M. Petry, 2010; Singer et al., 2020). In light of these similarities, the fifth edition of the “Diagnostic and Statistical Manual of Mental Disorders” categorizes gambling disorder in the category of “Substance-related and Addictive Disorders” (Association, 2013). In contrast to substance-use-disorders, differences in behavioral and/or neural effects between gamblers and controls are unlikely to be confounded by chronic or acute drug effects (Clark et al., 2019; Peters & Büchel, 2011; Wiehler & Peters, 2015). Gambling disorder has thus been termed a “pure addiction” (Mark Dixon, Ghezzi, et al., 2006).

Recently, categorical definitions of mental illness have increasingly been called into question. The National Institute for Mental Health of the United States proposed the Research Domain Criteria (RDoC) to foster characterization of the dimensions underlying psychiatric disorders. According to this approach, research in cognitive science should focus on the identification of continuous neuro-cognitive dimensions that might go awry in disease, i.e. trans-diagnostic markers (Nelson et al., 2016). Here we focus on two promising candidates for such trans-diagnostic processes that are affected across a range of psychiatric conditions, including gambling disorder: temporal discounting, i.e. the devaluation of delayed rewards (Bickel et al., 2019; Lempert et al., 2019; Peters & Büchel, 2011), and model-based (MB) control during reinforcement learning (Daw et al., 2011). MB control refers to computationally more expensive goal-directed strategies that utilize models of the environment, contrasting with model-free (MF) control that operates on stimulus-response associations (Balleine & O’Doherty, 2010; Daw et al., 2011; Doll et al., 2012; Valerie Voon et al., 2017).

Steep discounting has been consistently observed in substance use disorders and gambling disorder (Bickel et al., 2012; Bickel et al., 2019; MacKillop et al., 2011; Reynolds, 2006). Moreover, alterations in temporal discounting occur in a range of other disorders, including depression, bipolar disorder, schizophrenia and borderline personality disorder (Amlung et al., 2019), underlining the trans-diagnostic nature of this process. Changes in the contributions of MF and MB control have likewise been reported across multiple disorders, including gambling disorder (Wyckmans et al., 2019), schizophrenia (Culbreth et al., 2016), obsessive compulsive disorder (Gillan et al., 2020) and substance use disorders (Sebold et al., 2014). Reduced MB control is also reflected in sub-clinical psychiatric symptom severity (Gillan et al., 2016).

Addiction is known to be under substantial contextual control. Addiction-related cues and environments are powerful triggers of subjective craving, drug use and relapse. Incentive sensitization theory (T. Robinson & Berridge, 1993; Terry E. Robinson & Berridge, 2008) provides a theoretical framework that links such effects to a highly sensitized dopamine system that responds to drugs and addiction-related cues. Increased responses of the dopamine system to addiction-related cues (“cue-reactivity”) has been consistently observed in neuroimaging studies of human addicts (Courtney et al., 2016; Moeller & Paulus, 2018), and there is evidence that trans-diagnostic behavioral traits are likewise under contextual control. For example, regular gamblers discount delayed rewards substantially more steeply when tested in a gambling-related environment as compared to a neutral environment (Mark. Dixon, Jacobs, & Sanders, 2006). Similar effects have been observed in laboratory tasks that include gambling-related cues (Dale et al., 2019; Genauck et al., 2020; Miedl et al., 2014) but whether other putative trans-diagnostic traits such as MB control are under similar contextual control is unclear. Beyond, it is unclear whether gambling severity or maladaptive control beliefs (Raylu & Oei, 2004) modulate such effects.

Though rarely examined in naturalistic settings, contextual effects on trans-diagnostic dimensions of decision-making are of substantial clinical and scientific interest. Settings with high ecological validity might provide more informative insights into the central drivers of maladaptive behavior than laboratory-based studies (Anderson & Brown, 1984). If such trans-diagnostic traits are further exacerbated in e.g. addiction-related environments, this could constitute a mechanism underlying the maintenance and/or escalation of maladaptive behavior. Second, traits such as temporal discounting can be modulated (Bickel et al., 2011; Bickel et al., 2019; Lempert & Phelps, 2016) and could thus serve as a potential treatment target (Lempert et al. 2019).

The present pre-registered study thus had the following aims. First, we aimed to replicate the findings by Dixon et al. (Mark Dixon, Ghezzi, et al., 2006), who observed increased temporal discounting in gambling-related environments in regular gamblers, compared to neutral environments. Second, we extended their approach by including a modified version of the prominent 2-step sequential decision task (Daw et al., 2011) to test whether model-based control of behavior is likewise under contextual control. Reduced model-based control has been linked to a range of psychiatric conditions (see above) including gambling disorder (Wyckmans et al., 2019). Third, we directly tested for associations of contextual effects with gambling severity and working memory capacity. Finally, our tasks allowed for comprehensive computational modelling of choices and response time (RT) distributions. Analyses of reinforcement learning and decision-making have recently been shown to substantially benefit from an incorporation of RTs (Fontanesi et al., 2019; Pedersen et al., 2017; Peters & D’Esposito, 2020; Shahar et al., 2019; Wagner et al., 2020) via the application of sequential sampling models such as the drift diffusion model (DDM) (Forstmann et al., 2016). Such analyses yield additional insights into the latent processes underlying decision-making (Wagner et al., 2020) and can improve parameter stability (Shahar et al., 2019). To account for these recent developments, we complemented our pre-registered analyses with additional analyses of temporal discounting and reinforcement learning drift diffusion models (RLDDM).

## Methods

### Preregistration

This study was preregistered via the open science framework (https://osf.io/5ptz9/?view_only=62956a2afcd1495984db4be18c54b50a). We deviated from the pre-registered study design in the following ways. First, it was initially planned to use a lab-setting for the neutral (non-gambling) testing environment. However, this was changed following pre-registration to a café, which we felt was more similar to the gambling environment in terms of the presence of social cues and the overall level of distraction. Second, we initially aimed to include gamblers fulfilling at least one DSM-5 criterion for gambling disorder. This was adjusted to a stricter inclusion criterion of at least three DSM-5 criteria. Both of these changes were implemented before testing began.

To further account for recent developments in computational modelling we also complemented the pre-registered computational modeling analyses with additional analyses of RT distributions via temporal discounting and reinforcement learning DDMs (Fontanesi et al., 2019; Pedersen et al., 2017; Peters & D’Esposito, 2020; Wagner et al., 2020). As a model-free measure of intertemporal choice we used choice proportions of larger-but-later values instead of the area under the empirical discounting curve (AUC) (Myerson et al., 2001).

A-priori sample size was calculated based on results by Dixon et al. (2006) observed an effect size of *d* = .5 for the effect of gambling environments on temporal discounting in regular gamblers. Power analysis (Faul et al., 2007) yielded a minimum sample size of n = 26 with alpha error probability of .05 and power of .80. We then pre-registered a target sample size of n = 30.

### Participants

Participants were recruited via advertisements posted online and in local gambling venues. First, they were screened via a telephone interview to verify that they show evidence for problematic gambling behavior, with a primary gambling mode of electronic slot machines. Further inclusion criteria were age in the range of nineteen to fifty, no illegal drug use, and no history of neuropsychiatric disorders, current medication or a history of cardiovascular disease. The ethics committee of the University of Cologne Medical Center approved all study procedures.

Forty-two participants were then invited to a first appointment, were they provided written informed consent and completed a questionnaire assessment and a set of working memory tasks (see section on *background screening* below). Five participants dropped out during or after the first appointment. Four additional participants were excluded after the first appointment because they fulfilled less than three DSM-V criteria for gambling disorder. Two participants dropped out after the first experimental testing session, and one participant was excluded because he fell asleep twice during one testing session. Due to technical problems, we obtained complete datasets for thirty participants for the intertemporal choice task and twenty-nine participants for the two-step task, with twenty-eight participants overlapping.

### Overall procedure

Participants were invited to three appointments. At the first appointment (*baseline screening*; see below) participants were invited to our lab and performed a questionnaire assessment and four working memory tasks. Participants were randomly assigned to one of the two locations (café vs. casino) on the first experimental appointment (pseudorandomized location [first session neutral or gambling] and task-version; see section on tasks below). We label the café environment as neutral because no gambling associated cues were present. In both locations, the delay discounting task was completed first, followed by the 2-step task. Appointments were made on an individual basis but spaced within 7+-2 days and around the same time of day +-2 hours. The café environment was an ordinary café serving non-alcoholic drinks and snacks and furnished with 10 tables and approximately 50 m^2^ of size. Testing occurred while the café was in business as usual and experimenter and participant sat at a table next to a wall to assure some privacy. The café was usually moderately attended and testing occurred at the same spot for all participants, with only a few exceptions when this seat was taken. The gambling environment was a common slot-machine venue operated by a German gambling conglomerate. The experimenter and participant were seated at a table placed next to a wall in sight of the electronic gaming machines (EGMs). In total there were four EGMs in direct sight of the participant and a total of ten in the room (hidden by eye protection walls). The density of gambling related cues varied as a function of people playing at EGMs, background sounds e.g. sounds of winning or money dropping were all depended on regularly customers. However, in nearly all cases other people were playing EGMs in direct sight of the participants. The experimenter was granted permission to conduct research in two local gambling venues. Two chairs and a table to use for the experimental session were provided. In both locations, subjects were placed in such a way that neither experimenter nor customers could view their screen. Both tasks ran on a 15inch Laptop using the Psychopysics toolbox (Kleiner et al., 2007) running in Matlab (The MathWorks ©).

### Background screening

Participants filled out a battery of questionnaires regarding gambling related cognition (GRCS) (Raylu & Oei, 2004) and symptom severity (DSM-5;KFG,SOGS) (Falkai, 2015; Lesieur & Blume, 1987; J. Petry & Baulig, 1996), demographic evaluation and standard psychiatric diagnostic tools (see Supplemental Tables S1 and S2).

We assessed working memory capacity using a set of four working memory paradigms. First, in an Operation Span Task (Redick et al., 2012) subjects were required to memorize a sequence of letters while being distracted by math-operations. Second, in a Listening Span Task (adapted from the German version of the Reading Span Test developed by van den Noort et al. (van den Noort et al., 2008) subjects were required to listen to a series of sentences and had to recall the last word of each sentence. Last, subjects performed two different versions of a Digit Span Task (forward/backward) that were adopted from the Wechsler Adult Intelligence Scale (Wechsler, 2008). Here, participants listened to a series of numerical digits which they had to recall as a series in regular or reverse order. All working memory scores were *z*-transformed and averaged to obtain a single compound working memory score (*z*-score).

### Temporal discounting task

Participants performed 140 trials of a temporal discounting task where on each trial they made a choice between a smaller-but-sooner (SS) immediate reward, and a larger-but-later (LL) reward delivered after a specific delay. SS and LL rewards were randomly displayed on the left and right sides of the screen, and participants were free to make their choice at any time. While SS rewards were held constant at 20€. LL rewards were computed as multiples of the SS reward (task version 1: 1.05, 1.055, 1.15, 1.25, 1.35, 1.45, 1.55, 1.65, 1.85, 2.05, 2.25, 2.55, 2.85, 3.05, 3.45, 3.85; task version 2: 1.025, 1.08, 1.2, 1.20, 1.33, 1.47, 1.5, 1.70, 1.83, 2.07, 2.3, 2.5, 2.80, 3.10, 3.5, 3.80. Each LL reward from one version was then combined with each delay option for this version (in days): (either: 1, 7, 13, 31, 58, 122, or v: 2, 6 15, 29, 62, 118) yielding 140 trials in total. The mean larger LL magnitude was the same across task versions and the order was counterbalanced across subjects and session (neutral/gambling).

At the end of each session, one decision was randomly selected and paid out in the form of a gift certificate for a large online store, either immediately (in the case of an SS choice) or via email/text message after the respective delay (in the case of a LL choice).

### 2-step task

Participants performed a slightly modified version of the 2-step task, a sequential reinforcement-learning (Daw et al., 2011). Based on more recent suggestions (Kool et al., 2016) we modified the outcome stage by replacing the fluctuating reward probabilities (reward vs. no reward) with fluctuating reward magnitudes (Gaussian random walks with reflecting boundaries at 0 and 100, and standard deviation of 2.5). In total the task comprised 300 trials. Each trial consisted of two successive stages: In the 1^st^ stage (S1), participants chose between two fractals embedded in grey boxes. After taking an S1 action, participants transitioned to one of two possible 2^nd^ stages (S2) with fixed transition probabilities of 70% and 30%. In S2, participants chose between two new fractals each providing a reward outcome in points (between 0-100) that fluctuated over time. To achieve optimal performance, participants had to learn two aspects of the task. They had to learn the transition structure, that is, which S1 stimulus preferentially (70%) leads to which pair of S2 stimuli. Further, they had to infer the fluctuating reward magnitudes associated with each S2 stimulus..

In both versions, the tasks differed in the S1 and S2 stimuli, and in the fluctuating rewards in S2. However both task versions reward walks were equal in variance and mean and were presented in counterbalanced order per session (neutral/gambling). Participants were instructed about the task structure and performed 40 practice trials (with different random walks and symbols) at the first appointment (*Baseline screening*). Following task completion, points (*0.25) were converted to € and participants could win a bonus of up to 4.50€ that was added to the baseline reimbursement of 10€/h.

### Computational modeling and Statistical Analysis

#### Temporal discounting model

We applied a single-parameter hyperbolic discounting model to describe how subjective value changes as a function of LL reward height and delay (Mazur, 1987; Green and Myerson, 2004):

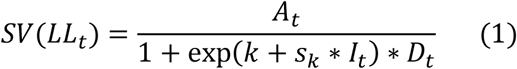

Here, *A*_*t*_ is the reward height of the LL option on trial *t, D*_*t*_ is the LL delay in days on trial *t* and *I*_*t*_ is an indicator variable that takes on a value of 1 for trials from the gambling context and 0 for trials from the neutral condition. The model has two free parameters: *k* is the hyperbolic discounting rate (modeled in log-space) and *s*_*k*_ is a weighting parameter that models the degree of change in discounting in the gambling compared with the neutral context condition.

#### Softmax action selection

Softmax action selection models choice probabilities as a sigmoid function of value differences (Sutton and Barto, 1998):

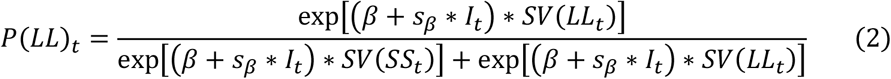

Here, *SV* is the subjective value of the larger but later reward according to Eq. 1 and *β* is an inverse temperature parameter, modeling choice stochasticity (for *β* = 0, choices are random and as *β* increases, choices become more dependent on the option values). *SV(SS*_*t*_) was fixed at at 20 and *I*_*t*_ is again the dummy-coded context regressor, and *s*_*β*_ models the context effect on *β*.

#### Temporal discounting drift diffusion models

To more comprehensively examine environmental effects on choice dynamics, we additionally replaced softmax action selection with a series of drift diffusion model (DDM)-based choice rules. In the DDM, choices arise from a noisy evidence accumulation process that terminates as soon as the accumulated evidence exceeds one of two response boundaries. In the present setting, the upper boundary was defined as selection of the LL option, whereas the lower boundary was defined as selection of the SS option.

RTs for choices of the SS option were multiplied by -1 prior to model fitting. We furthermore used a percentile-based cut-off, such that for each participant the fastest and slowest 2.5 percent of trials were excluded from the analysis. We then first examined a null model (DDM_0_) without any value modulation. Here, the RT on each trial *t* (*t* ϵ 1:140) is distributed according to the Wiener First Passage Time (*wfpt*):

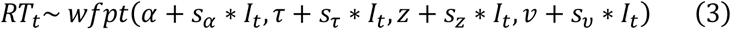

The parameter α models the boundary separation (i.e. the amount of evidence required before committing to a decision), τ models the non-decision time (i.e., components of the RT related to motor preparation and stimulus processing), *z* models the starting point of the evidence accumulation process (i.e., a bias towards one of the response boundaries, with *z*>.5 reflecting a bias towards the LL boundary, and *z*<.5 reflecting a bias towards the SS boundary) and v models the rate of evidence accumulation. Note that for each parameter *x*, we also include a parameter *s*_*x*_ that models the change in that parameter from the neutral context to the gambling context (coded via the dummy-coded condition regressor *I*_*t*_).

As in previous work (Pedersen et al., 2017; Fontanesi et al., 2019; Peters and D’Esposito, 2020, Wagner et al. 2020), we then set up temporal discounting diffusion models with modulation of drift rates by the difference in subjective values between choice options. First, we set up a version with linear modulation of drift-rates (DDM_lin_) (Pedersen et al., 2017):

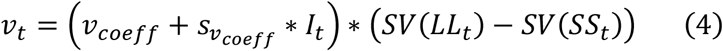

Here, the drift rate on trial *t* is calculated as the scaled value difference between the subjective LL and SS rewards. As noted above, RTs for SS options were multiplied by -1 prior to model estimation, such that this formulation predicts SS choices whenever SV(SS)>SV(LL) (the trial-wise drift rate is negative), and predicts longest RTs for trials with the highest decision-conflict (i.e., in the case of SV(SS)= SV(LL) the trial-wise drift rate is zero). We next examined a DDM with non-linear trial-wise drift rate scaling (DDM_S_) that has recently been reported to account for the value-dependency of RTs better than the DDM_lin_ (Fontanesi et al., 2019; Peters & D’Esposito, 2020; Wagner et al., 2020). In this model, the scaled value difference from Eq. 4 is additionally passed through a sigmoid function with asymptote *v*_*max*_:

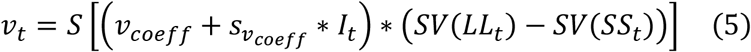

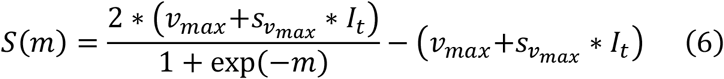

All parameters including *v*_*coeff*_ and *v*_*max*_ were again allowed to vary according to the context, such that we included *s*_*x*_ parameters for each parameter *x* that were multiplied with the dummy-coded condition predictor *I*_*t*_.

### Reinforcement Learning model

#### Hybrid model

We first applied a slightly modified version of the hybrid RL model (Daw et al., 2011) to analyze the strength of model-free and model-based RL strategies. The model updates MF state-action values (*Q*_*MF*_ -values, Eq. 7, 8) in both stages through prediction errors (Eq. 9, 10). In stage 1, MB state-action values (*Q*_*MB*_) are then computed from the transition and reward estimates using the Bellman Equation (Eq. 11).

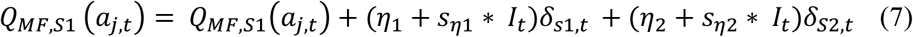

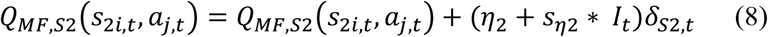

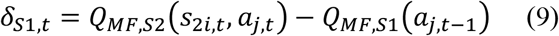

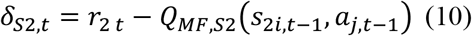

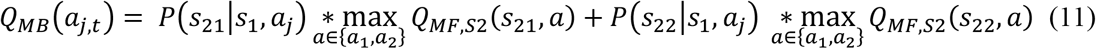

Here, *i* indexes the two different second stages (*S*_21_, *S*_22_), *j* indexes actions *a* (*a*_1_, *a*_2_) and *t* indexes the trials. Further, *η*_1_ and *η*_2_denote the learning rate for S1 and S2, respectively. S2 MF *Q*-values are updated by means of reward (*r*_2,*t*_)prediction errors (*δ*_*S*2,*t*_) (Eq. 8, 10). To model S1 MF *Q*-values we allow for reward prediction errors at the 2nd-stage to influence 1st-stage *Q*-values (Eq. 7, 9).

In addition, as proposed by Toyama et al. (Toyama et al., 2017, 2019) *Q*-values of all unchosen stimuli were assumed to decay with decay-rate η_decay_ and centered to the mean of reward walks (0.5). A decay of *Q*-values over time accounts for the fact that participants know that reward walks fluctuate over time. The decay was implemented according to Eq. 12 and 13:

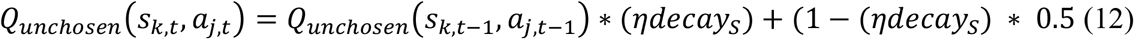

*where*

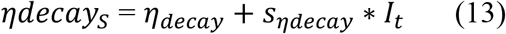

and *k ϵ* {1,21,22}, that is, *k* indexes the three task stages

S1 action selection is then modelled via weighting S1 MF and MB *Q*-values through a softmax action-selection. S2 stage action selection is likewise modelled as a function of MF *Q*-value differences. Separate ‘inverse temperature’ parameters *β* model subjects’ weights of MF and MB *Q*-Values (Eq. 14 and Eq. 15). The additional parameter *ρ* captures 1st-stage choice perseveration, and is set to 1 if the previous S1 choice was the same and is zero otherwise.

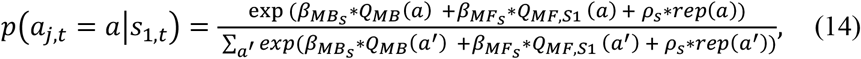

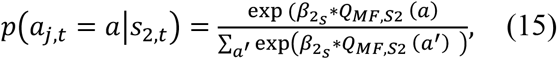

where:

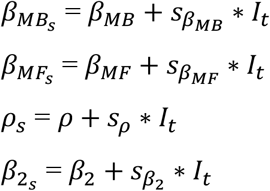

#### Hybrid model with drift diffusion action selection

As in our analysis of temporal discounting we replaced softmax action selection with a DDM choice rule (Shahar et al., 2019), leaving the reinforcement learning equations unchanged. For each stage of the task, the upper boundary was defined as selection of one stimulus, whereas the lower boundary was defined as selection of the other stimulus. We modelled each stage of the task using separate non-decision time (*τ*), boundary separation (α) and drift-rate (*v*) parameters. The bias (*z*) was fixed to 0.5. All parameters including *vcoeff*_*MF*_, *vcoeff*_*MB*_ and *v*_*max*_ were again allowed to vary according to the context, such that we included *s*_*x*_ parameters for each parameter *x* that were multiplied with the dummy-coded condition predictor *I*_*t*_ (see above). Data were filtered using a percentile-based cut-off, such that for each participant the fastest and slowest 2.5 percent of RTs/trials were excluded from further analysis. In addition, an absolute cutoff of > 150ms was applied. We then first examined a null model (DDM_0_; Eq. 3) without any value modulation followed by two value-informed models where the drift-rate (*v*) is a linear (Eq. 16 and 17) or sigmoid (Eq. 18) function of MF and MB *Q*-value weights. For the linear version, the drift rate in S1 is

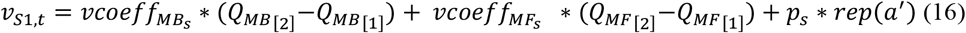

and the drift rate in S2 is calculated as

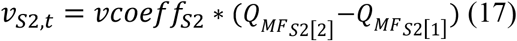

For the non-linear version, the linear drift rate from equations 16 and 17 are additionally passed through a sigmoid:

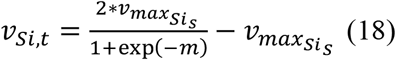

where

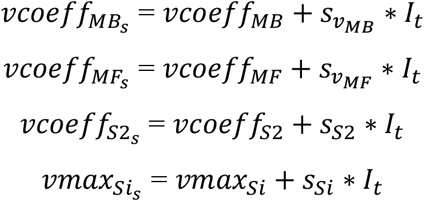

#### Hierarchical Bayesian models

Softmax models were fit to all trials from all participants using a hierarchical Bayesian modeling approach with separate group-level distributions for all baseline parameters for the neutral context and shift parameters (*s*_*x*_) for the gambling context.

For the intertemporal choice model fitting was performed using Markov Chain Monte Carlo (MCMC) sampling as implemented in the JAGS (Version 4.3) software package (Martyn Plummer, 2003) in combination with the Wiener module (Wabersich and Vandekerckhove, 2014). Model estimation was done in R (Version 4.0.3) using the corresponding R2Jags package (Version 0.6-1). For baseline group-level means, we used uniform priors defined over numerically plausible parameter ranges (see code and data availability section for details). For all *s*_*x*_ parameters modeling context effects on model parameters, we used Gaussian priors with means of 0. For group-level precisions, we used gamma distributed priors (.001, .001). We initially ran 2 chains with a varying burn-in period and thinning of two until convergence. Chain convergence was then assessed via the Gelman-Rubinstein convergence diagnostic 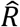 and sampling was continued until 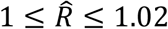 for all group-level and individual-subject parameters. 20k additional samples were then retained for further analysis.

For the 2-step task, model estimation was performed using MCMC sampling as implemented in STAN (Stan Development Team, 2020) via R (Version 4.0.3) and the rSTAN package (Version 2.21.0).

For baseline group-level means, we used uniform and normal priors defined over numerically plausible parameter ranges (see code and data availability section for details). For all *s*_*x*_ parameters modeling context effects on model parameters, we used Gaussian priors with means of 0. For group-level standard deviations we used cauchy (0, 2.5) distributed priors. We initially ran 2 chains with a burn-in period of 1000 and retained 2000 samples for further analysis. Chain convergence was then assessed via the Gelman-Rubinstein convergence diagnostic 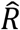 and sampling was continued until 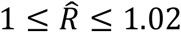. This threshold was not met for one participant 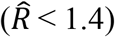.

For both tasks, relative model comparison was performed via the *loo*-package in R (Version 2.4.1) using the Widely-Applicable Information Criterion (WAIC) where lower values reflect a superior fit of the model (Vehtari et al., 2017). We then show posterior group distributions for all parameters of interest as well as their 85% and 95% highest density intervals. For group comparisons we report Bayes Factors for directional effects for *s*_*x*_ hyperparameter distributions of *s*_*x*_ > 0 (gambling context > neutral context), estimated via kernel density estimation using R via the RStudio (Version 1.3) interface. These are computed as the ratio of the integral of the posterior difference distribution from 0 to +∞ vs. the integral from 0 to -∞. Using common criteria (Beard et al. 2016), we considered Bayes Factors between 1 and 3 as anecdotal evidence, Bayes Factors above 3 as moderate evidence and Bayes Factors above 10 as strong evidence. Bayes Factors above 30 and 100 were considered as very strong and extreme evidence respectively, whereas the inverse of these reflect evidence in favor of the opposite hypothesis.

#### Posterior Predictive checks

We carried out posterior predictive checks to examine whether models reproduced key patterns in the data, in particular the value-dependency of RTs (Peters & D’Esposito, 2020; Wagner et al., 2020) and participant’s choices. For the intertemporal choice task, we binned trials of each individual participant into five bins, according to the absolute difference in subjective larger-later vs. smaller-sooner value (“decision conflict”, computed according to each participant’s median posterior log(k) parameter from the DDM_S_, and separately for the neutral and gambling context. For each participant and context, we then plotted the mean observed RTs as a function of decision conflict, as well as the mean RTs across 10k data sets simulated from the posterior distributions of the DDM_0_, DDM_lin_ and DDM_S_. For the 2-step task, we extracted mean posterior parameter estimates and simulated 1000 datasets in R (Version 4.0.3) using the Rwiener package (Version 1.3.3). We then show RTs as a function of S2 reward difference of observed data and the mean RTs across 1000 simulated datasets for of all DDMs. We further show that our models capture the relationship of S2 reward differences and optimal (max[reward]) choices.

#### Model free analysis

As a model-agnostic measure of temporal discounting, we examined arcsine-square-root transformed proportions of LL choices as a function context (neutral vs. gambling) with order (neutral vs. gambling session first) as fixed and subject as random effect using a hierarchical generalized linear model (HGLM). For the 2-step task we likewise use a HGLM approach and modeled 2nd-stage RTs as a function of transition (common vs. rare) and context (neutral vs. gambling) as fixed and subject as random effect. In line with our modelling analyses, data were filtered so that implausibly fast RTs were excluded (see Methods). A standard analysis of stay probabilities (Daw et al., 2011) adapted to our task version is reported in the Supplement (Supplemental Table S5).

#### Subjective Craving Rating

On each testing day, participants rated their subjective craving (“How much do you desire to gamble right now?”) on a visual-analogue scale ranging from 0 to 100, both at the beginning of the testing session, and at the end following task completion. We then used paired t-tests to examine whether subjective craving differed between the testing environments (neutral vs. gambling).

## Results

### Subjective craving

Craving was assessed on a visual-analogue-scale before and after task performance. Due to technical problems, ratings of the first eight participants were lost. Another two participants did not complete post-task ratings. In the remaining n=22 participants, craving was substantially higher in the gambling-related environment compared to the neutral environment (paired t-test pre-task: t_23_ = -3.13; p = 0.0048, Cohen’s *d*: 0.75; post-task: t_21_ = -4.32, p = 0.0003, Cohen’s *d* = 0.68; see Figure 1).

**Figure 1.**
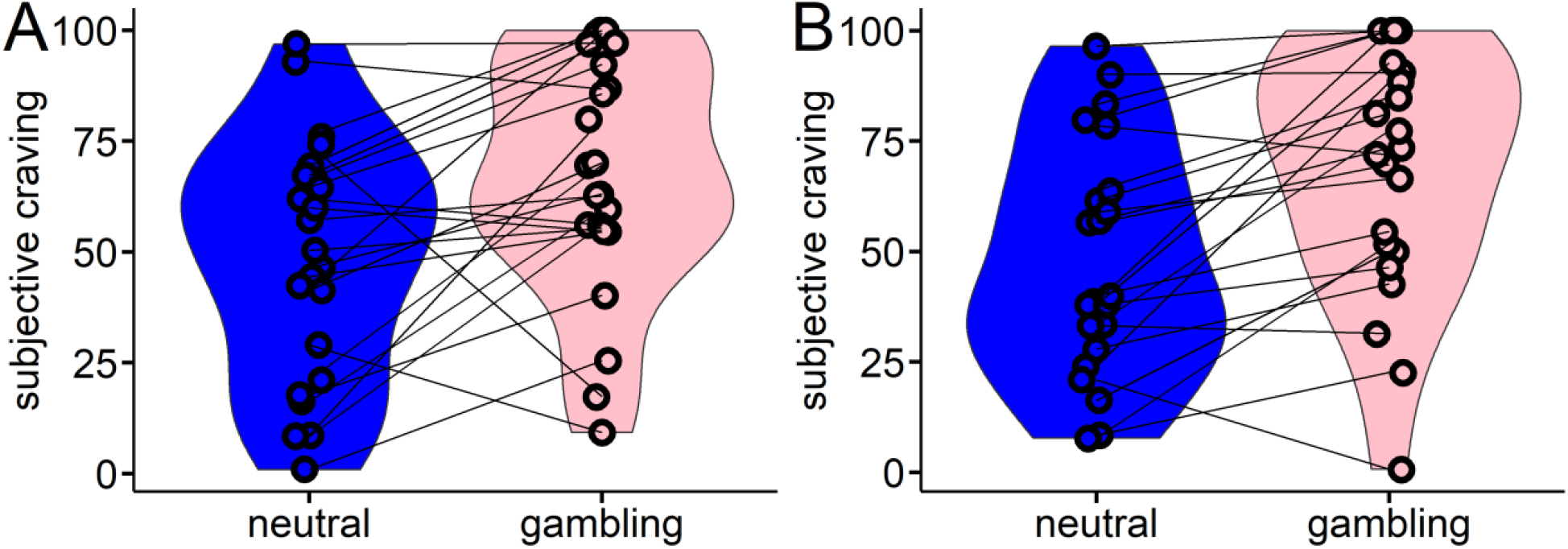
Subjective craving was assessed at the beginning (A) and at the end (B) of each testing session via a visual-analogue scale rating. Craving was significantly higher in the gambling environment, both at the start of the session (p = 0.0048) and at the end of the session (p = 0.0003).

### Temporal discounting

#### Model-agnostic analysis temporal discounting task

Raw proportions of larger-but-later (LL) choices are plotted in Figure 2A for each context. A hierarchical linear model on arcsine-square-root transformed proportion values with context (gambling vs. neutral) and order (gambling first vs. neutral first) as fixed effects and subject as random effect confirmed a significant main effect of context (F_28_ = 13.33, p = 0.01) such that participants made more LL selections in the neutral vs. the gambling-related environment. There was no effect of order on choice proportions. Overall response time (RT) distributions are plotted in Figure 2B with choices of the LL option coded as positive RTs and choices of the smaller-sooner option coded as negative RTs.

**Figure 2.**
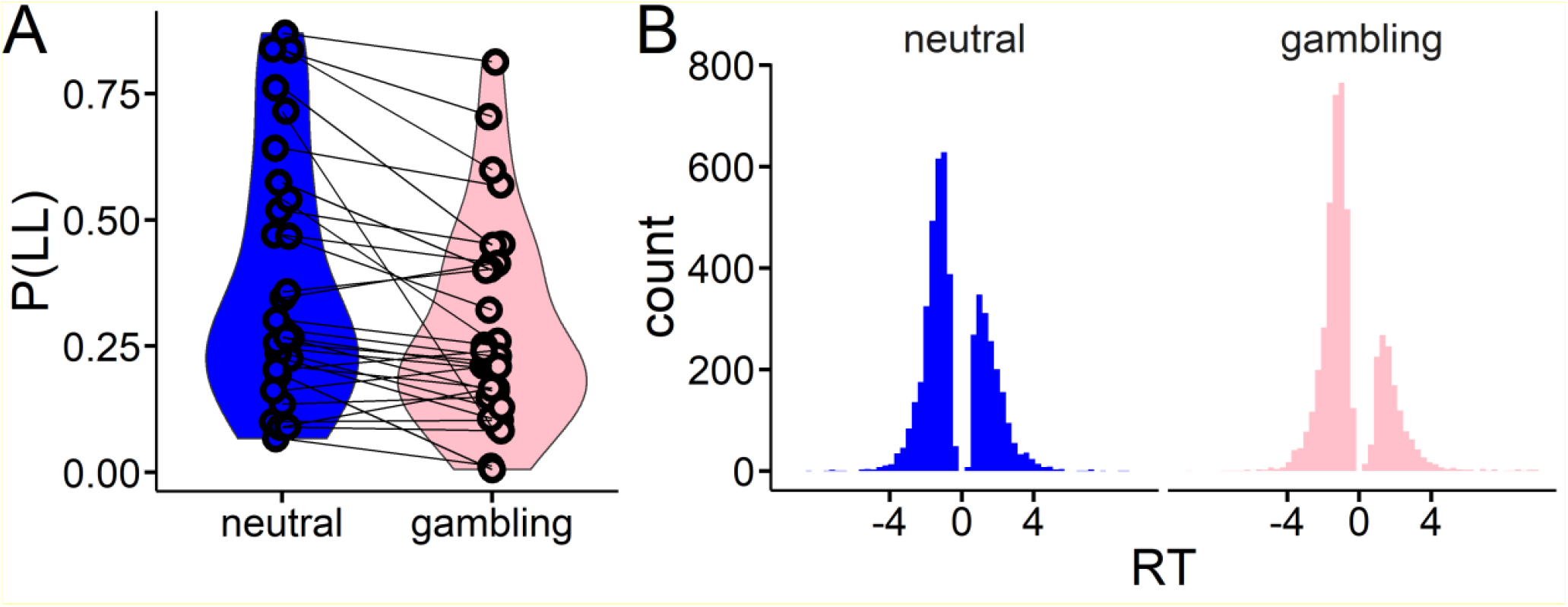
Behavioral data from the temporal discounting task. A: raw proportions of larger-later (LL) choices in each context. B: Overall response time distributions with choices of the LL option coded as positive RTs and choices of the smaller-sooner option coded as negative RTs; Note, this was done to add choice coding to the computational model.

#### Softmax choice rule

We first modeled the data using standard softmax action selection. This analysis revealed an overall context effect on log(k), such that discounting was substantially steeper in the gambling context compared to the neutral context (Figure 3B, 95% HDI > 0). Examination of Bayes Factors indicated that an increase in log(k) in the gambling context (s_k_) was about 116 times more likely than a decrease (see Figure 3 and Table 3). There was no evidence for a change in choice stochasticity (softmax[β]; Figure 3C/D).

**Figure 3.**
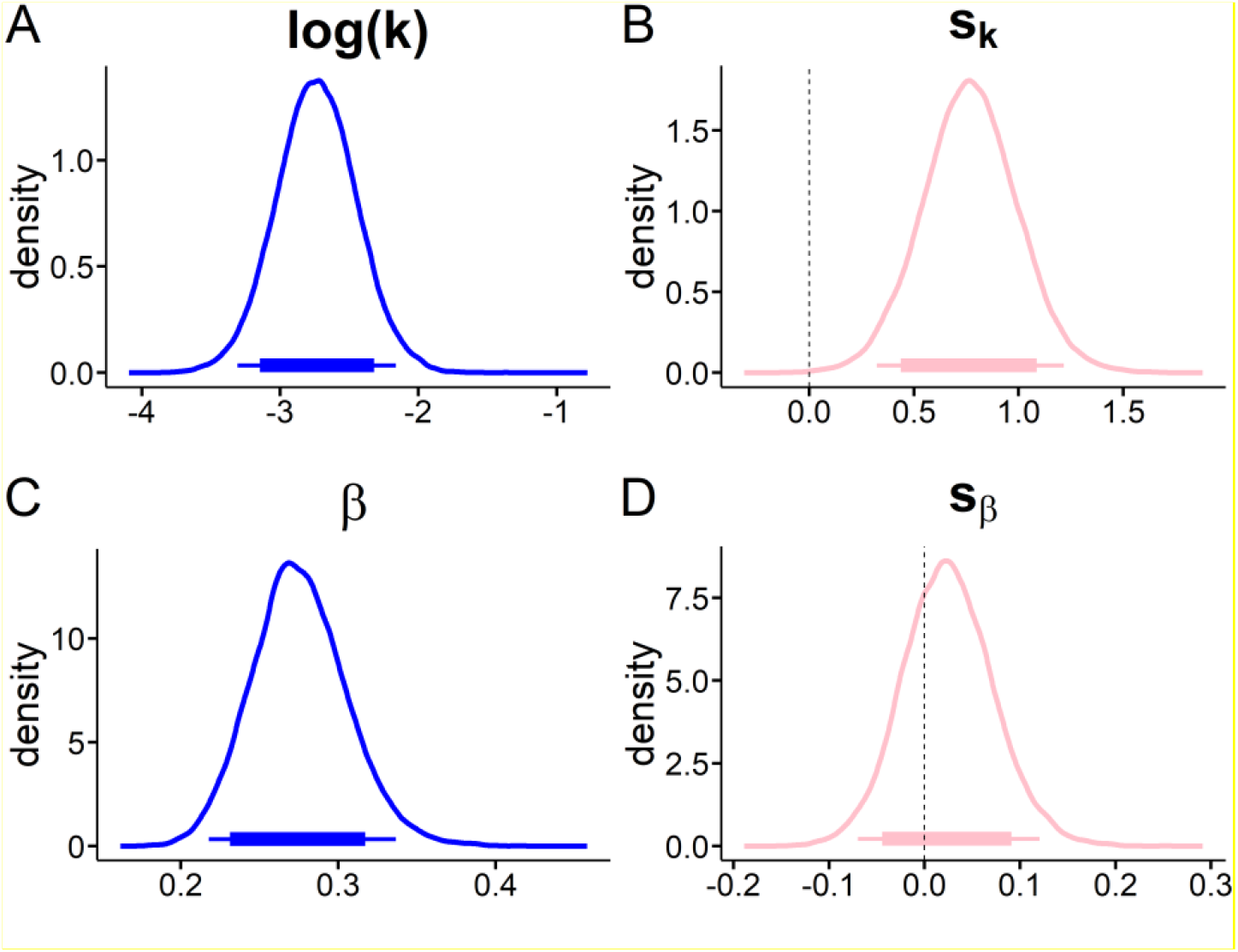
Softmax model; Posterior distributions of mean hyperparameter distributions for the neutral baseline context (blue) and the corresponding shift in the gambling context (pink). A, discount-rate log (k); B, shift in discount-rate (s_k_); C, softmax β; D, shift in softmax β; Thin (thick) horizontal line denote 95% (85%) highest posterior density intervals

#### Temporal discounting drift diffusion models (DDMs)

Model comparison of temporal discounting DDMs revealed the same model ranking in each context (Supplemental Table S3) such that the data were best accounted for by a temporal discounting DDM with non-linear drift rate scaling. This model accounted for around 90% of decisions (Supplemental Table S4, Supplemental Figure S1) and posterior predictive checks confirmed that it reproduced individual-participant RTs (Supplemental Figure S2).

We next examined the posterior distributions of model parameters of the best-fitting TD-DDM model (DDMs with sigmoid drift rate scaling; we further report model comparison, binary choice predictions and posterior predictive checks in the corresponding *Model comparison and validation* section in the supplement). Results are plotted in Figure 4 and Figure 5 and Bayes Factors for all context-effects are listed in Table 1. There was a consistent positive association between trial-wise drift rates and value differences in the neutral context (Figure 4E, the 95% HDI for the drift rate coefficient parameter did not include 0). Likewise, there was a numerical bias towards the smaller-sooner option in the baseline condition (85% HDI < 0.5, see Figure 4F). The non-decision time was numerically smaller in the gambling context (85 % HDI < 0, Figure 5B, Table 1), amounting to on average a 50ms faster non-decision time. The maximum drift-rate was substantially higher in the gambling context (95% HDI > 0, Figure 5D).

**Figure 4.**
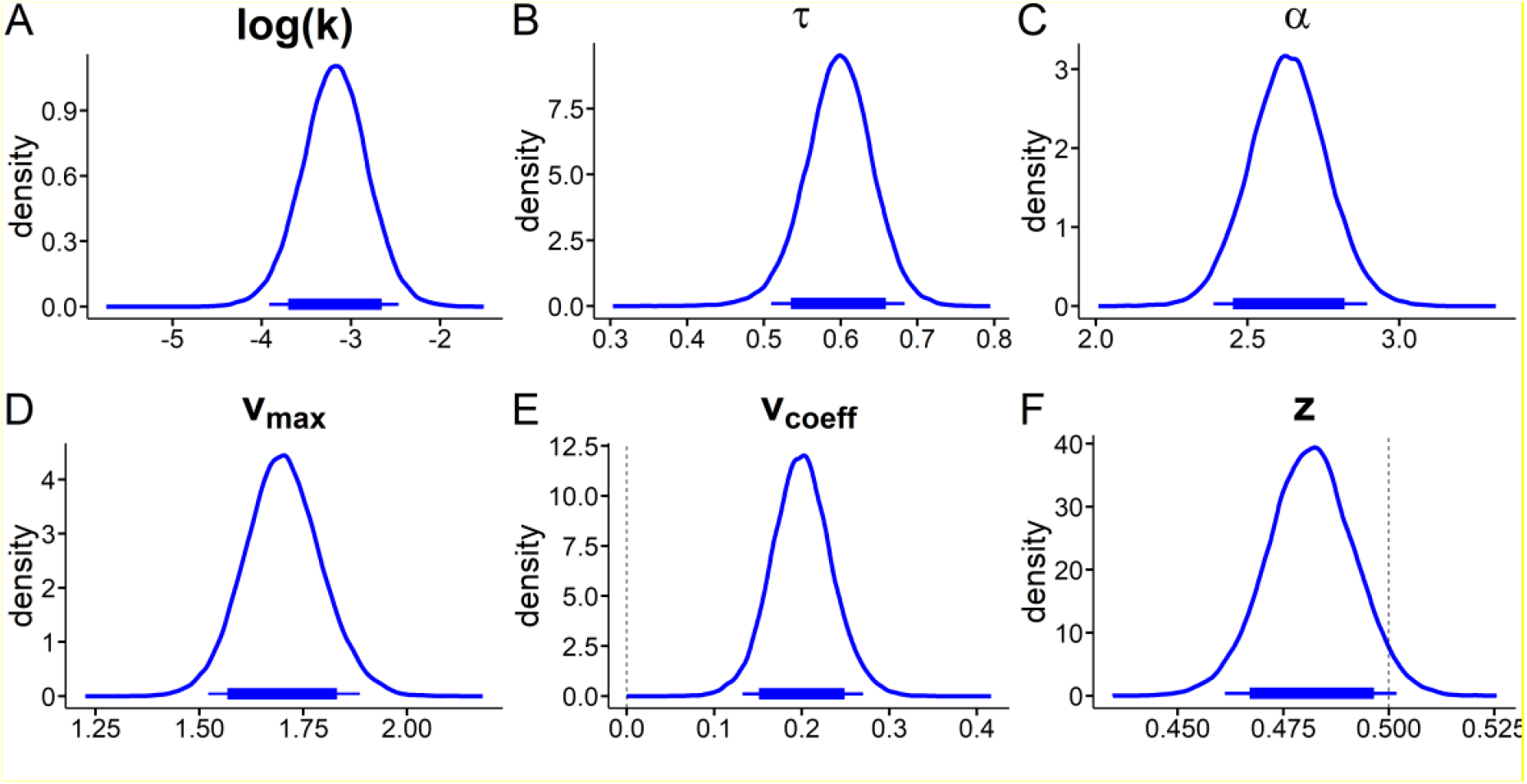
Temporal discounting drift diffusion model results: posterior distributions for hyperparameter means from the neutral context. A: discount-rate log(k), B: non-decision time τ, C: boundary separation α, D: maximum drift-rate v_max_, E: drift-rate coefficient v_coeff_, F: starting-point *z*. Thin (thick) horizontal line denote 95% (85%) highest posterior density intervals

**Figure 5.**
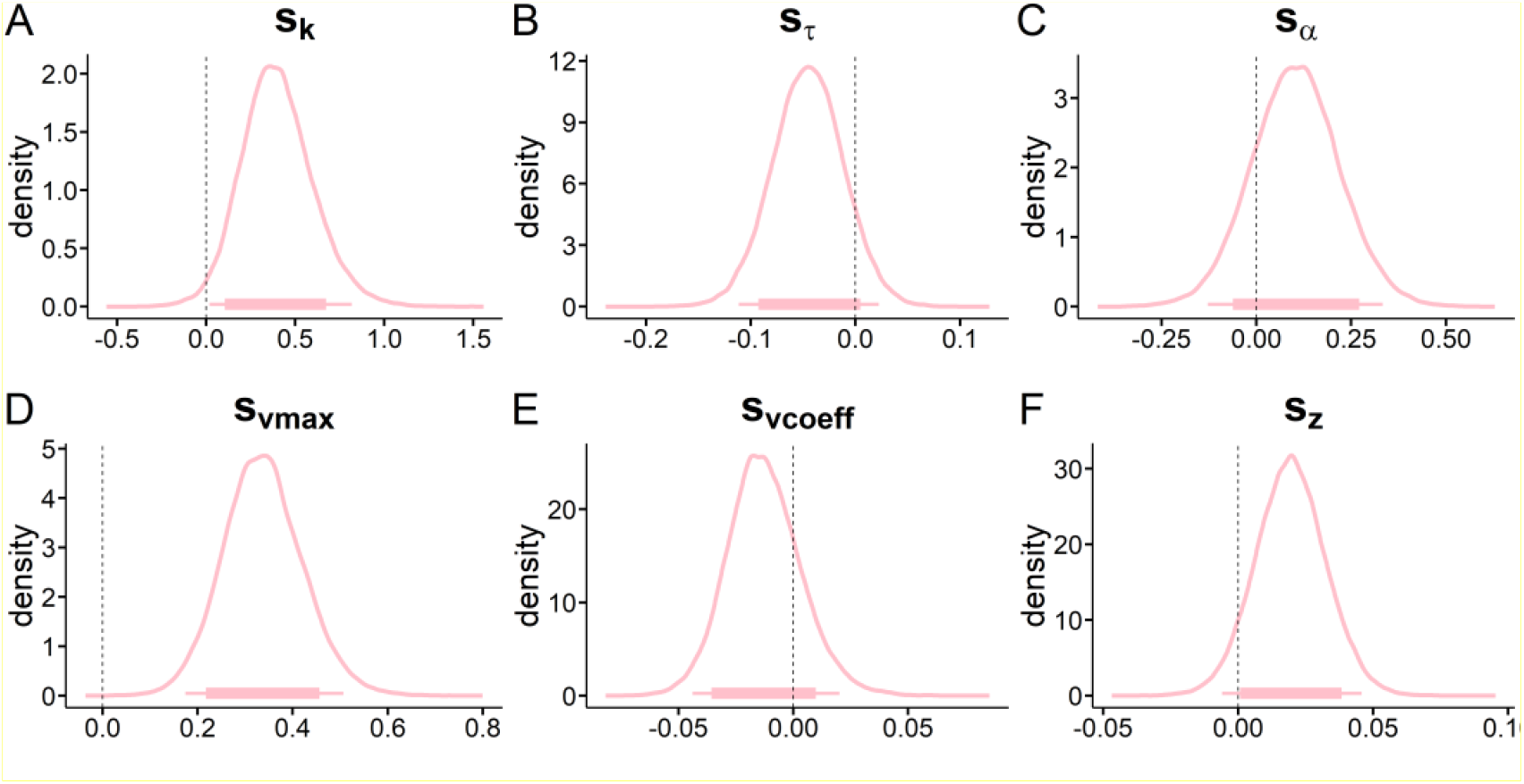
Temporal discounting drift diffusion model results: posterior distributions for hyperparameter means for context shift (s_x_) parameters modeling changes from the neutral to the gambling context. A: shift in discount-rate (s_k_), B: shift in non-decision time s_τ_, C: shift in boundary separation s_α_, D: shift in maximum drift-rate v_max_, E: shift in drift-rate coefficient v_coeff_, F: shift in starting-point s_*z*_. Thin (thick) horizontal line denote 95% (85%) highest posterior density intervals.

**Table 1.** Overview of overall context differences. For group comparisons we report Bayes Factors for directional effects for s_x_ hyperparameter distributions of s_x_ > 0 (gambling context > neutral context).

| Model parameter<br>(change in gambling context) | Softmax Model |  | DDM<sub>s</sub> |  |
| --- | --- | --- | --- | --- |
|  | *Mean* | *dBf* | *Mean* | *dBf* |
| $s_k$ (discount-rate) | 0.77 | 1688.53 | 0.40 | 54.20 |
| $s_\beta$ (softmax beta) | 0.025 | 2.27 | - | - |
| $s_{vcoeff}$ (drift-rate coeff.) | - | - | -0.012 | 0.25 |
| $s_\tau$ (non-decision time) | - | - | -0.05 | 0.10 |
| $s_a$ (boundary separation) | - | - | 0.10 | 4.40 |
| $s_z$ (starting point bias) | - | - | 0.02 | 13.64 |
| $s_{vmax}$ (max drift-rate) | - | - | 0.33 | 39490.71 |

As in the softmax model (Figure 3), we observed a substantial increase in the discount rates log(k) in the gambling context (95% HDI > 0, see Figure 5A, Table 1).

#### Temporal discounting and gambling-related questionnaire data

As preregistered, we next examined whether the increased in discount-rate s_k_ in the gambling context was associated with symptom severity or gambling related cognition. We therefore computed a compound symptom severity *z*-score of DSM-5 (Falkai, 2015), SOGS (Lesieur & Blume, 1987) and KFG (J. Petry & Baulig, 1996) scores. Gambling context-related changes in temporal discounting were not significantly associated with symptom severity (rho = -0.05, p = 0.78) but were positively associated with Gambling Related Cognition Scale (Raylu & Oei, 2004) (rho = 0.39; p = 0.03); see Figure 6). There were no significant correlations between changes in craving and changes in discounting or working memory capacity and temporal discounting (Supplemental Results 1).

**Figure 6.**
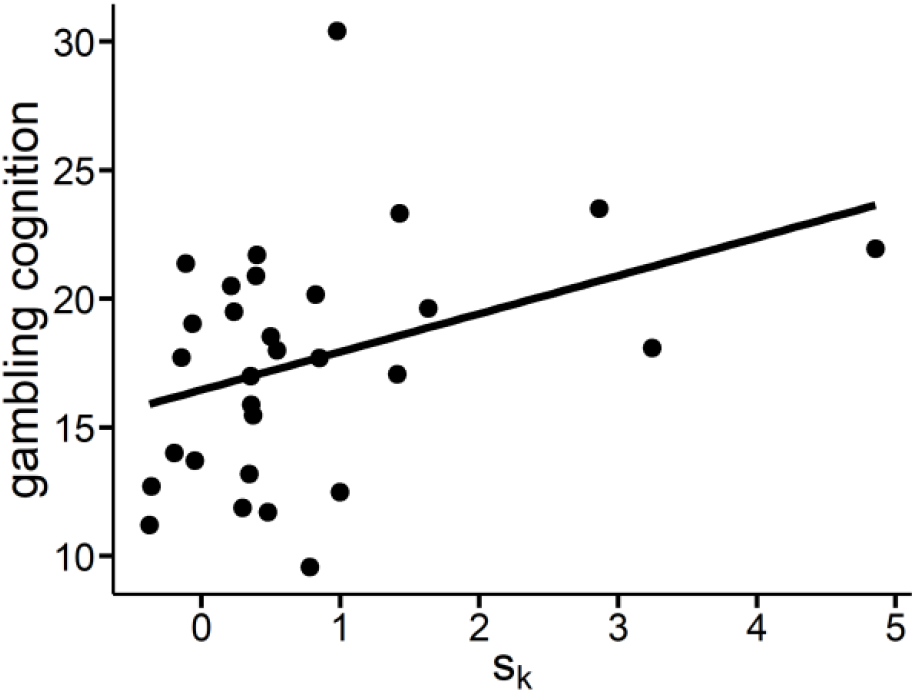
Relationship of total scores from the gambling-related cognition scale (GRCS) (Raylu & Oei, 2004) and changes in discount-rate from neutral to gambling environment (s_k_)[softmax model].

### 2-step reinforcement learning task

#### Model-agnostic analysis 2-step task

Participants earned significantly more points in the gambling context (t-test: t_28_ = -2.44, *p* = 0.02, Cohen’s *d* = 0.22). For S2 RTs, we observed a significant main effect of transition (Supplemental Table S7 and Supplemental Figure S3) and a trend for a transition x context interaction (p = 0.07; see Supplemental Table S7), reflecting increased model-based control (Otto et al., 2015; Shahar et al., 2019).

An analysis of stay probabilities adapted to the present 2-step task version is shown in Supplemental Table S5. In each context, we observed main effects of reward (reflecting model-free RL) and reward x transition interaction (reflecting model-based RL). The reward x transition x context interaction was not significant.

#### Hybrid model with softmax choice rule

We first examined a hybrid model (Daw et al., 2011) with extensions by ourselves and Otto et al. (Otto et al., 2015) using a standard softmax choice rule (see Methods; Figure 7). This model included separate parameters for S1 and S2 learning rates, model-free and model-based β weights for S1 and a β weight for S2 *Q*-value differences. We confirmed substantial contributions of both MB and MF values to S1 choices (Figure 7B,C). There was an increase in the S2 learning-rate η (95% HDI > 0, Figure 7F) in the gambling context. Furthermore, there was a strong decrease in MF β weights (95% HDI < 0, Figure 7H) such that participants showed substantially less MF behavior in the gambling environment compared to the neutral environment. BFs for directional effects indicate that an increase in MB reinforcement learning is 4 times more likely than a decrease. For examination of Bayes Factors see Table 2.

**Table 2.** Overview of overall context differences. For context comparisons we report Bayes Factors or directional effects for s_x_ hyperparameter distributions of s_x_ > 0 (gambling context > neutral context).

| Model parameter (shift) | Softmax Model |  | DDMs |  |
| --- | --- | --- | --- | --- |
|  | *Mean* | *dBF* | *Mean* | *dBF* |
| $s_{\eta S1}$ (learning-rate S1) | 0.44 | 3.29 | 0.0801 | 1.186 |
| $s_{\eta S2}$ (learning-rate S2) | 0.40 | 92.3 | 0.280 | 14.658 |
| $s_{\tau S1}$ (non-decision times S1) | - | - | 0.001 | 0.8454 |
| $s_{\tau S2}$ (non-decision times S2) | - | - | 0.001 | 1.161 |
| $s_p$ (Stickiness S1) | 0.04 | 1.946 | 0.05 | 2.365 |
| $s_{a S1}$ (boundary separation S1) | - | - | -0.002 | 0.9354 |
| $s_{a S2}$ (boundary separation S2) | - | - | 0.0149 | 2.026 |
| $\beta_{MF} / S_{vcoeffMF}$ (MF beta/ drift-rate coeff.) | -1.14 | 0.010 | -0.93 | 0.083 |
| $\beta_{MB} / S_{vcoeffMB}$ (MB beta/ drift-rate coeff.) | 1.08 | 4.00 | 4.01 | 169.62 |
| $\beta_{S2} / S_{vcoeffS2}$ (S2 beta / drift-rate coeff.) | -0.44 | 0.428 | -0.64 | 0.271 |
| $s_{vmax S1}$ (max drift-rate S1) | - | - | -0.19 | 0.296 |
| $s_{vmax S2}$ (max drift-rate S2) | - | - | 0.41 | 15.83 |

**Figure 7.**
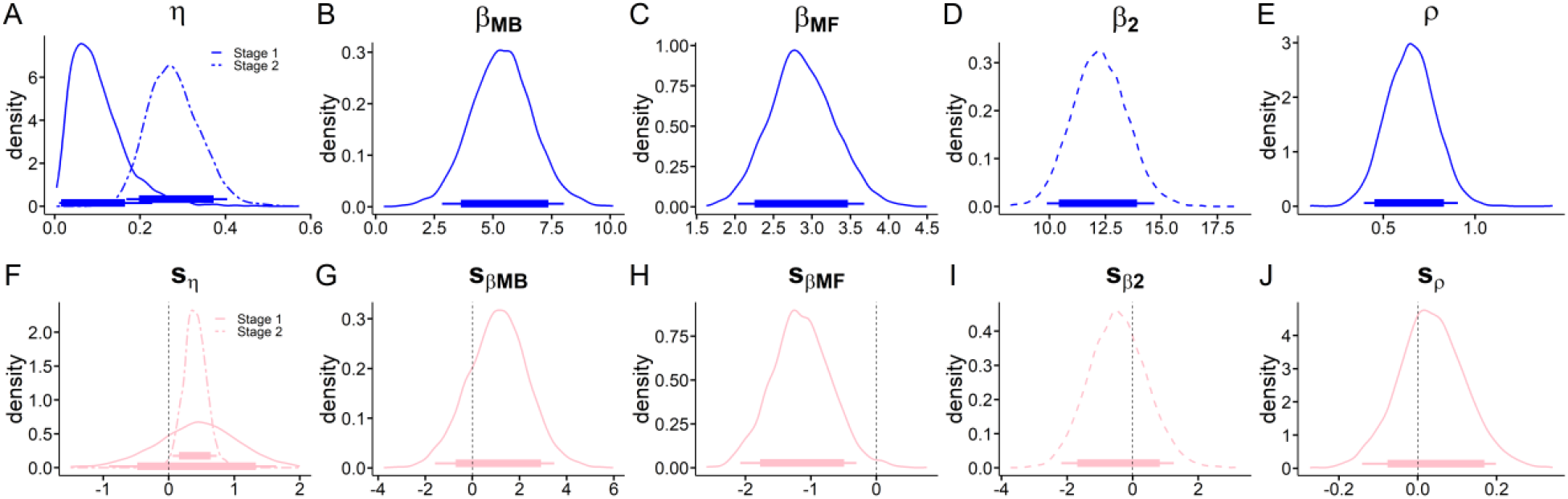
Hybrid model with softmax choice rule posterior distributions (top row: neutral context, bottom row: parameter changes in gambling context) of all group level means. A, S1 and S2 learning-rates. B, MB β weight. C, MF β weight. D, S2 β weight. E, perseveration parameter ρ.F, shift in S1 and S2 learning rates. G, shift in MB β. H, shift in MF β. I, shift in S2 β. J, shift in stickiness parameter ρ. Thin (thick) horizontal line denote 95% (85%) highest posterior density intervals.

#### Hybrid model with drift diffusion choice rule

We next combined the hybrid model with a DDM choice-rule (Shahar et al., 2019) and likewise compared DDMs that varied in the way that they accounted for the influence of *Q*-value differences on trial-wise drift rates in both task stages. Model comparison yielded the same model ranking in each context, such that the data were best accounted for by an RLDDM with non-linear drift rate scaling (Supplemental Table S8). This model accounted for around 73% of S1 choices, and around 81% of S2 choices (Supplemental Table S9). Posterior predictive checks confirmed that this model reproduced the observed RTs (Supplemental Figure S4) and choice proportions (Supplemental Figure S5).

Posterior distributions for the best-fitting RLDDM are shown in Figure 8 (neutral context parameters) and Figure 9 (gambling context changes). We observed positive associations between trial-wise drift rates and *Q*-value differences (Figure 8F-J, all 95% HDIs above 0). Likewise, as in the softmax model, beta weights were positive, indicating MB and MF effects on behavior (Figure 8E-G, all 95% HDIs > 0). In the gambling context, we observed a decrease in the MF component (85% HDI < 0) and a robust increase in MB contributions (95% HDI > 0). BFs for directional effects are provided in Table 2. Overall, these results suggest decreased MF and increased MB reinforcement learning due to gambling context exposure.

**Figure 8.**
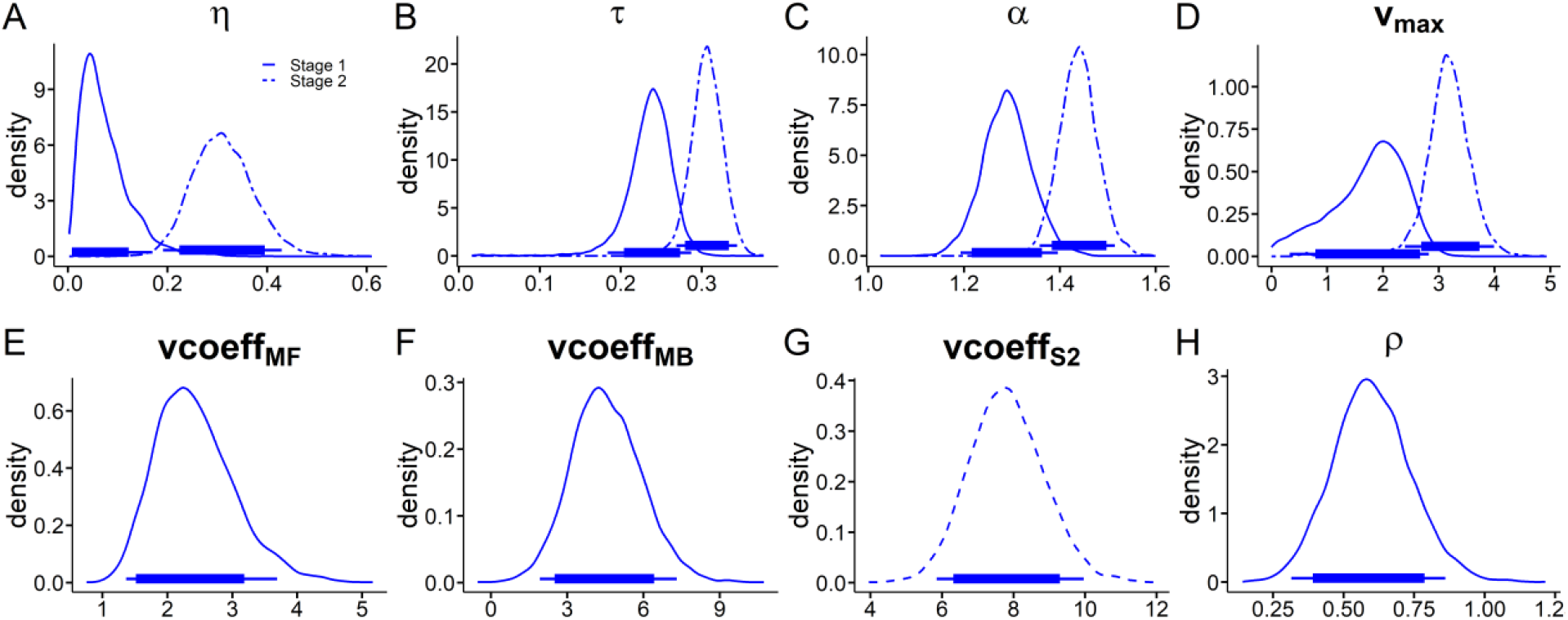
RL-DDM. Posterior distributions of all hyperparameters for the neutral baseline condition. A: S1 and S2 learning rates η. B: S1 and S2 non-decision time τ. C: S1 and S2 boundary separation α. D: S1 and S2 drift-rate maximum v_max_. E: MF drift-rate coefficient vcoeff_MF_. F: MB drift-rate coefficient vcoeff_MB_. G: S2 drift-rate coefficient vcoeff_S2_. H: stickiness parameter ρ. Thin (thick) horizontal line denote 95% (85%) highest posterior density intervals.

**Figure 9.**
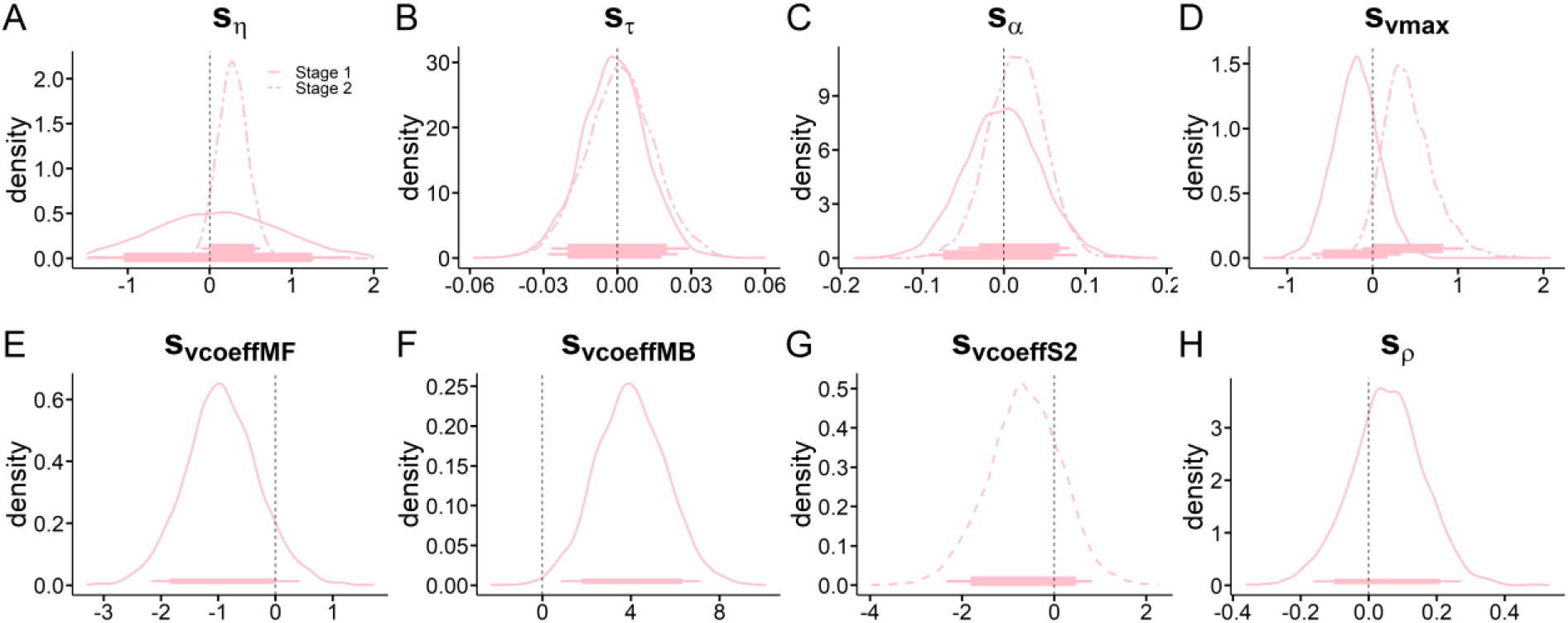
RL-DDM. Posterior distributions of all hyperparameters shift-parameters modelling the change from neutral to gambling condition. A, shift in Stage 1 and Stage 2 learning rates η. B, shift in S1 and S2 non-decision time τ. C, shift in S1 and S2 boundary separation α. D, shift in S1 and S2 drift-rate maximum v_max_. E, shift in S1 MF drift-rate coefficient vcoeff_MF_. F, shift in S1 MB drift-rate coefficient vcoeff_MB_. G, shift in S2 drift-rate coefficient vcoeff_S2_. H, shift in stickiness parameter ρ. Thin (thick) horizontal line denote 95% (85%) highest posterior density intervals.

#### Reinforcement learning and gambling related questionnaire data

As preregistered, we examined associations between ρ (perseveration/stickyness) and gambling severity (average *z*-score across SOGS (Lesieur & Blume, 1987), KFG (J. Petry & Baulig, 1996) and DSM-5 criteria). The association was non-significant ρ (r = -0.10, p = 0.59). There were no significant correlations between changes in craving and changes in MB behavior, nor between MB behavior and working memory capacity (Supplemental Results 2).

## Discussion

Here we comprehensively examined the contextual modulation of two putatively trans-diagnostic markers implicated in addiction, temporal discounting (Bickel et al., 2019; Lempert et al., 2019) and model-based control (Gillan et al., 2016; Gillan et al., 2020) in a pre-registered study. We studied regular slot machine gamblers, a group previously characterized by high levels of temporal discounting (Wiehler & Peters, 2015) and reduced model-based control (Wyckmans et al., 2019). Following a seminal study by Dixon et al. (Mark. Dixon, Jacobs, & Sanders, 2006), regular gamblers were tested in gambling environments (slot-machine venues) and neutral control environments. Gambling cue exposure modulated temporal discounting and model-based control in gamblers in opposite ways: replicating Dixon et al., (2006), discounting substantially increased in a gambling context. In contrast, model-based (MB) control improved (increased). This differential modulation of two prominent trans-diagnostic traits in (behavioral) addiction has important theoretical and clinical implications.

Theoretical accounts highlight the central role of addiction-related cues and environments in drug addiction (T. Robinson & Berridge, 1993). Similar mechanisms have been suggested to underlie gambling disorder (M. J. F. Robinson et al., 2016). Because terrestrial slot machine gambling is directly linked to specific locations, gambling disorder is uniquely suited to investigate the impact of cue exposure on behavior. We replicated the finding of Dixon et al. (2006) of steeper discounting in gambling vs. neutral environments in gamblers. This effect was observed across model agnostic analyses (proportion of LL choices) and computational modeling (softmax, drift diffusion models [DDM]). We additionally extended these earlier results in the following ways. First, we observed an association of this effect with maladaptive control beliefs (GRCS) (Raylu & Oei, 2004) suggesting that such beliefs contribute to increased temporal discounting in gambling environments. Second, in a subset of participants, we confirmed that exposure to gambling environments substantially increases subjective craving. Third, comprehensive modeling via DDMs revealed additional effects on latent decision processes. The gambling context-related attenuation in non-decision time mirrors previous effects of pharmacological enhancement of dopamine transmission (Wagner et al. 2020). In contrast to these earlier pharmacological results, we observed a substantial *increase* in maximum drift rate (*V*_*max*_) in the gambling context, reflecting increased value sensitivity of RTs. Lastly, our results complement cue-reactivity designs showing increased impulsive and/or risky choice in gamblers during exposure to gambling cues in laboratory studies (Dale et al., 2019; Genauck et al., 2020; Miedl et al., 2014). However, effect sizes during naturalistic cue exposure (e.g. the present study and Dixon et al., 2006) were substantially larger than during lab-based exposure in these previous studies.

In addition to temporal discounting, we included a 2-step sequential decision-making task designed to dissociate model-based (MB) from model-free (MF) contributions to behavior (Daw et al., 2011). Reductions in MB control are associated with compulsivity-related disorders (Gillan et al., 2016; Gillan et al., 2020; V. Voon et al., 2015b). We observed increased MB learning and reduced MF learning in gamblers in the gambling context, a pattern of results consistent between softmax and DDM models. These findings were again corroborated by model-agnostic analyses. First, participants earned more points in the gambling context, an effect linked to MB learning (Kool et al., 2016). Second, the slowing of RTs following rare transitions, an indirect measure for MB learning (Otto et al., 2015) tended to be more pronounced in the gambling vs. neutral context. The MF effect correlated with gambling severity in an exploratory analysis, such that higher gambling severity was associated with a greater reduction in MF reinforcement learning in the gambling context. Together, these findings converge on the picture of *decreased* MF and *increased* MB control in gamblers when tested in gambling-related environments.

The latter result contrast with our pre-registered hypothesis of *reduced* MB control, which was based on findings of reduced MB control in populations with extensive habit formation (Gillan et al., 2016; Gillan et al., 2020; V. Voon et al., 2015a). Addiction is likewise thought to be inherently associated with pathological habits (Barry J Everitt & Trevor W Robbins, 2005; Robbins & Everitt, 1999) which are thought to be triggered by exposure to environmental cues (Antons et al., 2020). We thus hypothesized gambling environments would likewise trigger increased MF behavior and reduced MB behavior on the 2-step task. However, critics of habit theory have emphasized that addiction might in contrast be associated with excessive goal-directed behavior, in particular in the presence of addiction-related cues (Hogarth 2020). Our findings are more in line with this latter perspective. This interpretation is compatible with incentive sensitization theory (T. Robinson & Berridge, 1993; Terry E. Robinson & Berridge, 2008), which proposes that addiction-related environments exert their influence on behavior in part via a potentiation in dopamine release (Anselme & Robinson, 2013; Berridge, 2016; T. E. Robinson & Berridge, 2001). Earlier studies observed increased MB control following increases in DA neurotransmission (Sharp et al., 2016; Wunderlich et al., 2012), which could contribute to the present findings regarding 2-step task performance. Furthermore, our results are compatible with decreased MF control under L-Dopa (Kroemer et al., 2019). The gambling context might thus enhance goal-directed control via an improved construction and/or utilization of the task transition structure. This interpretation further resonates with other perspectives on DA function including a regulation of outcome sensitivity or precision (FitzGerald et al., 2015; Shiner et al., 2012), or the general motivation to exert (cognitive) effort (Berke, 2018). The observed increase in S2 learning rates could likewise be mediated in part by increases in DA transmission (Frank & O’Reilly, 2006).

If the effects of gambling environments on 2-step task performance are (at least in part) driven by increases in DA, then the question arises why gamblers at the same time exhibited substantially increased temporal discounting. The literature on DA effects on temporal discounting is a mixed bag (D’Amour-Horvat & Leyton, 2014) with some studies showing reduced discounting (van Gaalen et al., 2006; Wagner et al., 2020), some increased discounting (Pine et al., 2010) and others suggesting baseline-dependent effects (Petzold et al., 2019).

Given that DA was neither measured nor directly manipulated here, these issues cannot be directly resolved. However, our data might nonetheless provide some insights. Effects of DA on decision-making might depend on both task and context (Mikhael et al., 2021). Under this view, DA signals average reward in the environment (context) and its effects on performance further differ as a function of task controllability [see (Mikhael et al., 2021) for details]. DA might thus facilitate cognitive control (Ott & Nieder, 2019; Westbrook et al., 2020) when cognitive effort requirements are high, and there is control over the outcome (e.g. 2-step task). In contrast, DA might facilitate impulsive choice for cognitively less demanding tasks (e.g. temporal discounting task) that are performed in an addiction-related context (Antons et al., 2020; Terry E. Robinson & Berridge, 2008) signaling high reward (Mikhael et al., 2021). A further mechanism known to modulate temporal discounting is episodic future thinking or future prospection (Gershman & Bhui, 2020; Peters & Büchel, 2010). Future prospection has been shown to attenuate temporal discounting in a range of settings (Rösch et al., 2021) and might be attenuated at gambling venues. Participants might be generally focused on the present in the presence of cues or contexts endowed with high levels of incentive salience (Flagel et al., 2009).

Our results show that two prominent (potentially trans-diagnostic) computational processes, temporal discounting and MB control, are differentially modulated by addiction-related environments in regular slot machine gamblers. This provides a computational psychiatry perspective on factors that contribute to the understanding of this disorder. The substantial contextual effects on temporal discounting further highlight the potential clinical relevance of this process (Amlung et al., 2019; Lempert et al., 2019). Gambling disorder is reliably associated with increased temporal discounting (Mark. Dixon et al., 2003; Mark. Dixon, Jacobs, & Sanders, 2006; MacKillop et al., 2011; Miedl et al., 2012; Wiehler & Peters, 2015). This trait-like behavior then appears to be further exacerbated during exposure to gambling-related environments, potentially contributing to the maintenance of maladaptive behavior. In contrast, MB control improved (increased) in a gambling context, despite the fact that an earlier study reported reduced MB control in gamblers (Wyckmans et al., 2019). In general these findings are further compatible with a greater tendency for pattern matching (Wilke et al., 2014) or enhanced cause-effect associations that might translate into increased MB control (Joukhador et al., 2004) and studies suggesting that DA increases the willingness to spend cognitive effort (Westbrook et al., 2020; Westbrook et al., 2021). 2-step task transitions are not random, but can be learned and exploited. An increased tendency to seek for patterns during gambling context exposure might facilitate this behavior. Our findings suggest that gamblers do generally show MB control, which contrasts in parts with one recent study (Wyckmans et al., 2019). This is supported by the robust RTs increases observed following rare transitions (Supplemental Table S7, Supplemental Figure S3) and the positive MB parameters observed across models, somewhat contrasting with the findings of Wyckmans et al. (Wyckmans et al., 2019), although different 2-step task versions have been used in these studies.

We also extended previous studies on this topic via a recent class of value-based decision models based on the DDM (Fontanesi et al., 2019; Pedersen et al., 2017; Peters & D’Esposito, 2020; Shahar et al., 2019; Wagner et al., 2020). Comprehensive RT-based analysis revealed that standard DDM parameters were largely unaffected by context, suggesting that primarily MF and MB contributions to evidence accumulation were affected by gambling environments (Figure 10.). Posterior predictive checks showed that a DDM with non-linear trial-wise drift rate scaling captured the relationship of decision conflict (SS-LL value difference) and RTs, replicating prior findings (Peters & D’Esposito, 2020; Wagner et al., 2020). We previously reported good parameter recovery of such temporal discounting DDMs (Peters & D’Esposito, 2020; Wagner et al., 2020).

A number of limitations need to be acknowledged. First, as in the original study (Mark. Dixon, Jacobs, & Sanders, 2006) we did not test a non-gambling control group. However, the observed associations between experimental effects and gambling severity / gambling-related cognition suggests that these effects are at least in part driven by the underlying problem gambling symptoms. Second, MB and MF effects in the 2-step task might be affected by instructions (da Silva & Hare, 2019). Participants in our study were well instructed in written and verbal form and completed extensive training trials. Moreover, due to the counter-balanced within-subject design, the observed context-dependent changes in MB/MF behavior cannot be attributed to overall instruction effects. Third, MB control might more generally be related to attentional or motivational processes. For example, incentives boost 2-step task performance (Patzelt et al., 2019). However, we ensured that mean and variance of reward walks as well as incentives were identical in both contexts. Fourth, although participants were tested in the same venues, the number of customers present varied across participants, affecting e.g. noise levels and auditory gambling cues (slot machine sounds etc.). A trade-off between the control of such variables and ecological validity is unavoidable when testing in naturalistic settings. Finally, DA neurotransmission was obviously not assessed, rendering our interpretation of the effects in terms of the incentive sensitization theory speculative. But the substantial increase in subjective craving supports the idea that cue exposure had subjective effects predicted by incentive sensitization.

To conclude, here we show that two computational trans-diagnostic markers with high relevance for gambling disorder in particular and addiction more generally are modulated in opposite ways by exposure to real gambling environments. Gamblers showed increased temporal discounting in a gambling context, and this effect was modulated by maladaptive control beliefs. In contrast, MB control improved, a finding that posits a challenge for habit/compulsion theories of addiction. Ecologically valid testing settings such as those investigated here can thus yield novel insights into environmental drivers of maladaptive behavior underlying mental disorders.

## Acknowledgements

This work was supported by Deutsche Forschungsgemeinschaft (PE1627/5-1 to JP)

## Author contributions

JP conceived the idea and acquired the funding. JP and BJW designed the study. BJW acquired the data. BJW analyzed the data and performed the modeling. DM contributed analytical tools/software. BJW wrote the first draft of the paper. BJW and JP wrote the paper. DM provided revisions. JP supervised the project.

## Supplemental Information

**Supplemental Table S1.**
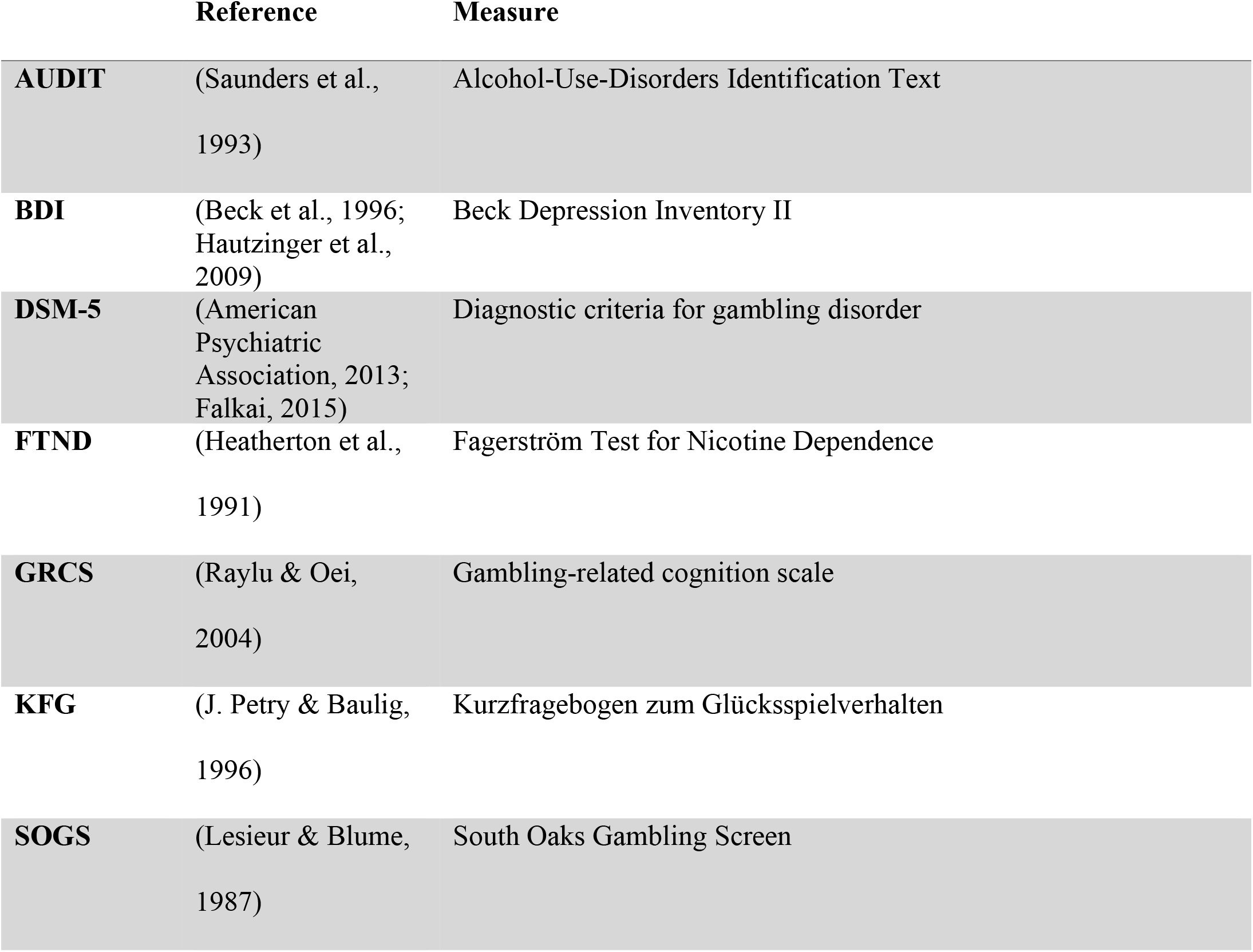
Baseline screening questionnaires.

|  | Reference | Measure |
| --- | --- | --- |
| AUDIT | (Saunders et al., 1993) | Alcohol-Use-Disorders Identification Text |
| BDI | (Beck et al., 1996; Hautzinger et al., 2009) | Beck Depression Inventory II |
| DSM-5 | (American Psychiatric Association, 2013; Falkai, 2015) | Diagnostic criteria for gambling disorder |
| FTND | (Heatherton et al., 1991) | Fagerström Test for Nicotine Dependence |
| GRCS | (Raylu & Oei, 2004) | Gambling-related cognition scale |
| KFG | (J. Petry & Baulig, 1996) | Kurzfragebogen zum Glücksspielverhalten |
| SOGS | (Lesieur & Blume, 1987) | South Oaks Gambling Screen |

**Supplemental Table S2.**
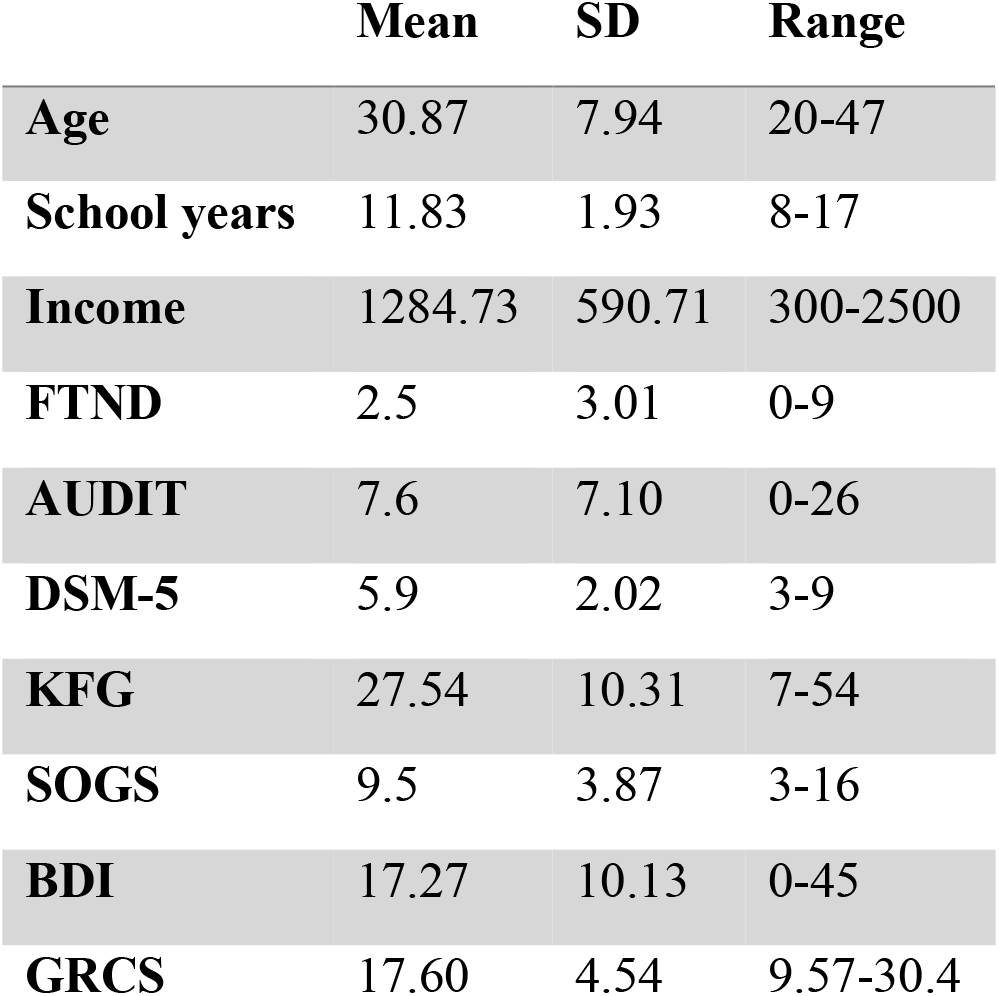
Summary of demographics and clinical information (n = 30).

|  | Mean | SD | Range |
| --- | --- | --- | --- |
| **Age** | 30.87 | 7.94 | 20-47 |
| **School years** | 11.83 | 1.93 | 8-17 |
| **Income** | 1284.73 | 590.71 | 300-2500 |
| **FTND** | 2.5 | 3.01 | 0-9 |
| **AUDIT** | 7.6 | 7.10 | 0-26 |
| **DSM-5** | 5.9 | 2.02 | 3-9 |
| **KFG** | 27.54 | 10.31 | 7-54 |
| **SOGS** | 9.5 | 3.87 | 3-16 |
| **BDI** | 17.27 | 10.13 | 0-45 |
| **GRCS** | 17.60 | 4.54 | 9.57-30.4 |

### Temporal discounting drift diffusion models (DDMs)

#### Model comparison and validation

We compared three versions of the drift diffusion model (DDM) that varied in the way that they accounted for the influence of value differences on trial-wise drift rates, based on model-fit (WAIC). To verify comparable model ranking across conditions, we first carried out a model comparison separately for each environment (see Supplemental Table S3). In both environments, a DDM with nonlinear drift-rate scaling (DDM_s_) (Fontanesi et al., 2019; Peters & D’Esposito, 2020; Wagner et al., 2020) accounted for the data best when compared to a DDM with linear scaling (DDM_lin_) (Pedersen et al., 2017) and a null model without value modulation (DDM_0_).

We then build a full model with group level distributions for the baseline condition (neutral context) and s_x_ parameters for each model parameter x, modeling the change from the neutral to the gambling context. S_x_ parameters where modeled with Gaussian priors with means of zero (see methods section). Model ranking was confirmed for the full model (Supplemental Table S3). We next compared the DDMs and the softmax model with respect to the proportion of binary choices (LL vs. SS selections) that they correctly accounted for. As can be seen from Supplemental Table S4, the DDM_S_ and DDM_lin_ performed numerically on par with the softmax model, whereas the DDM_0_ performed substantially worse (see Supplemental Figure S1, Supplemental Table S4). Posterior predictive checks for the winning model showed that it accurately captured the effect of decision conflict (value difference) on RTs (see section *Posterior Predictive Checks* below and Supplemental Figure S2). Parameter recovery for this model was reported in our prior papers (Peters & D’Esposito, 2020; Wagner et al., 2020).

**Supplemental Table S3.** Temporal discounting DDM model comparison using the Watanabe-Akaike Information Criterion (WAIC) revealed the same model ranking for each context (neutral vs. gambling) and the full model. Scores are WAIC (SE).

|  | Neutral | Gambling | Full model |
| --- | --- | --- | --- |
| **DDM<sub>0</sub>** | 12037.7 (150.1) | 11754.9 (157.7) | 23792.4 (217.4) |
| **DDM<sub>lin</sub>** | 9304.6 (155.5) | 9174.7 (158.5) | 18949.6 (219.0) |
| **DDM<sub>s</sub>** | 8982.3 (155.6) | 8744.6 (157.6) | 17656.0 (220.8) |

**Supplemental Table S4.** Proportions of correctly predicted binary choices (mean [range]) for the temporal discounting models (neutral vs. gambling context; see Supplemental Figure S1).

|  | Neutral | Gambling |
| --- | --- | --- |
| **Softmax** | 0.89 [0.57-0.98] | 0.90 [0.66-1.00] |
| **DDM<sub>0</sub>** | 0.74 [0.52-0.93] | 0.76 [0.55-0.99] |
| **DDM<sub>lin</sub>** | 0.89 [0.58-0.98] | 0.90 [0.65-0.99] |
| **DDM<sub>s</sub>** | 0.90 [0.60-0.99] | 0.91 [0.70-1.00] |

**Supplemental Figure S1.**
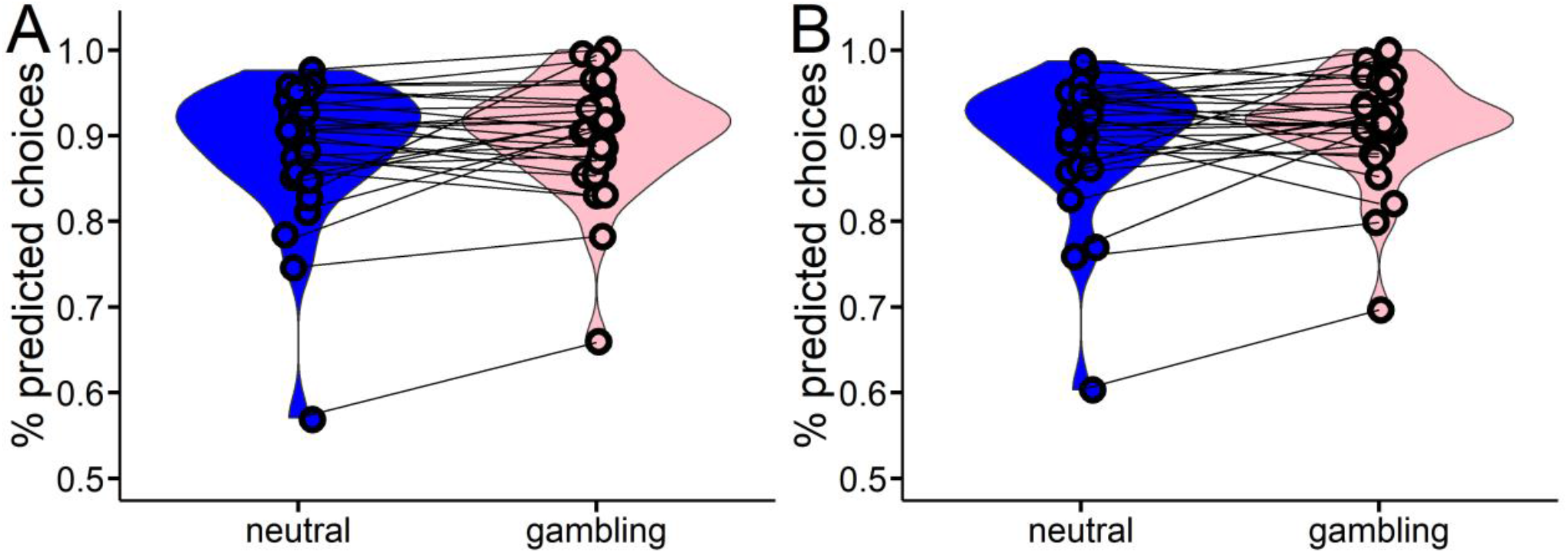
Proportions of correctly predicted binary choices for the softmax model (A) and the drift diffusion model with non-linear drift rate scaling (B, DDMs) in both contexts (neutral [blue], gambling [pink]).

### Posterior predictive checks

#### Temporal discounting

We carried out posterior predictive checks to visualize if our computational analysis captures key aspect in the data, in particular the value-dependency of RTs (Peters & D’Esposito, 2020; Wagner et al., 2020). For the temporal discounting task, we binned trials per participant into five bins according to the absolute difference in larger-later vs. smaller-sooner value (“decision conflict”, computed according to each participant’s median posterior log(k) parameter from the DDM_S_, and separately for the neutral and gambling context conditions). We then plotted the mean observed RTs as a function of decision conflict per participant and context, as well as the mean RTs across 10.000 data sets simulated from the posterior distributions of the DDM_0_, DDM_lin_ and DDM_S_ (see Supplemental Figure S2).

**Supplemental Figure S2.**
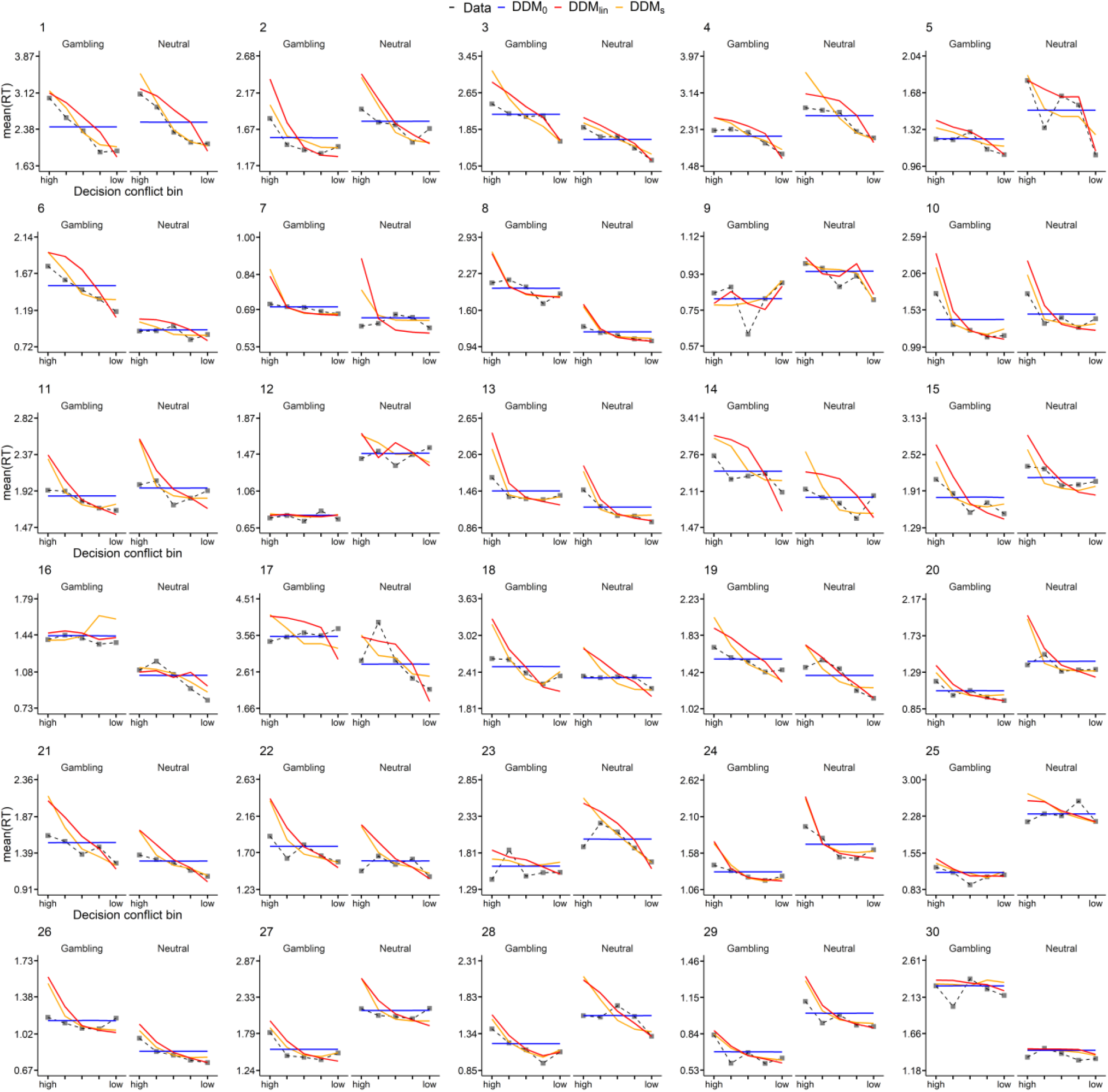
Posterior predictive checks for temporal discounting drift diffusion models. For each participant and condition (Gambling vs. Neutral), trials were binned into five equal sized bins according to the absolute difference between subjective LL and SS option values (decision conflict bin). Plotted are mean observed RTs per bin (data) as well model-generated RTs (blue: DDM_0_, red: DDM_lin_, orange: DDM_S_) averaged over 10,000 datasets simulated from the respective posterior distributions of the hierarchical models.

**Supplemental Table S5.**
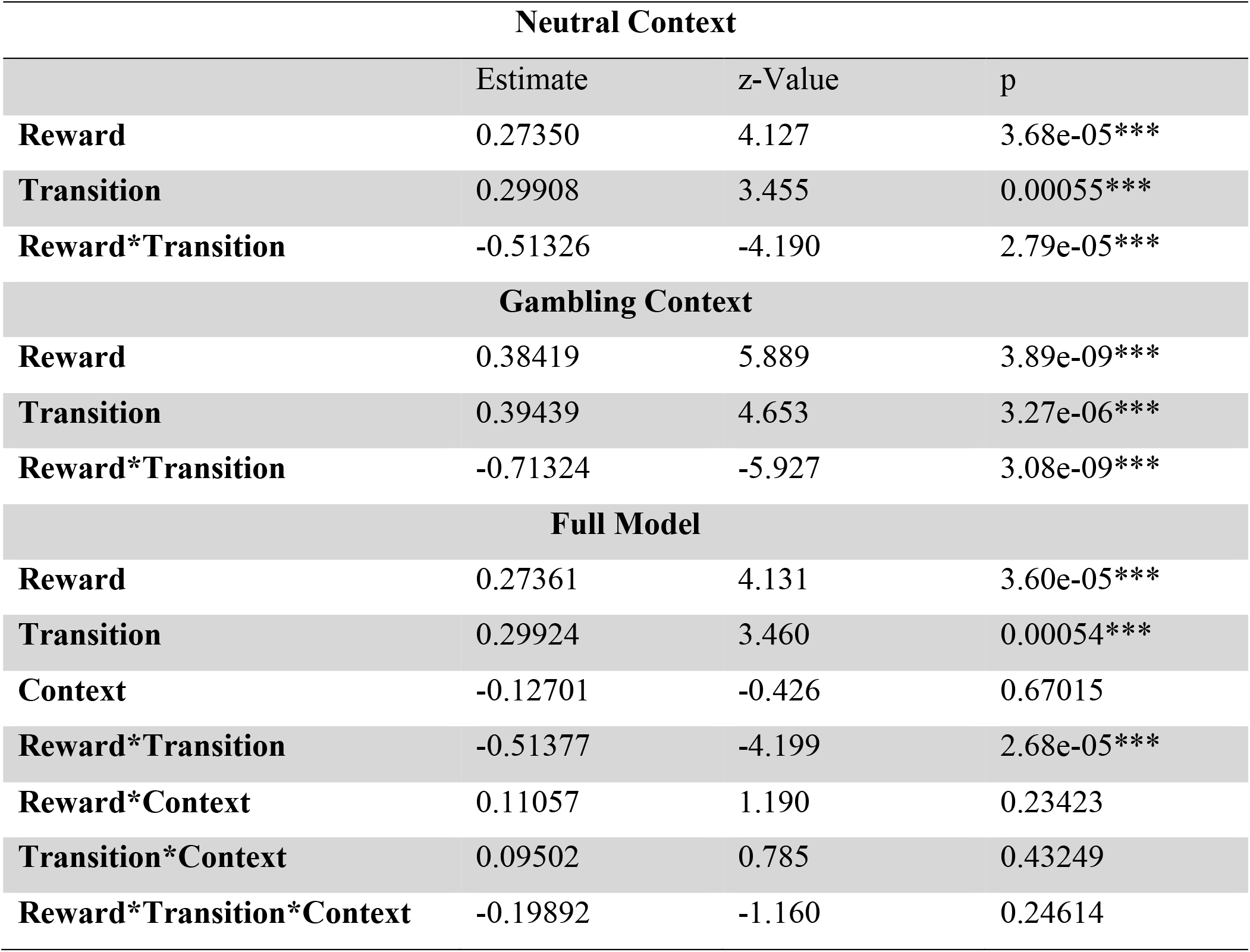
Model agnostic analysis of stay probability via a hierarchical general linear model (HGLM). HGLMs were estimated for each context separately using reward and transition as fixed and subject as random effects. The full model with stay probability as dependent variable included the predictors reward, transition (rare vs. common) and context (gambling vs. neutral) as fixed effects and subject as random effect. As a model-agnostic performance measure, the probability of choosing the same S1 option as in the previous trial (stay-probability) is typically analyzed as a function of reward, transition, and their interaction (Daw et al., 2011). Since the 2-step task version employed here utilized continuous payoffs, every trial was rewarded. The “reward” in S2 can thus not be used to directly predict stay probabilities, as done in previous work. Therefore, here the “reward” factor is computed relative to a moving average of recent rewards as a reference. Specifically, we categorized a reward R_t-1_ as positive “R+” if R_t-1_ was higher than the mean of last 7 rewards (R_t_ > mean[R_t-1: t-7_]) and as negative “R-” if R_t_ < mean(R_t-1:t-7_).

#### Model free analysis of Stage 1 RTs

S1 RTs were modeled as a function of categorized reward in the previous trial (see previous section for how this was defined) and context as fixed effects and trial and subject as random effects. Previous reward significantly increased RTs (t = -2.431, p = 0.015, see Supplemental Table S6). We also observed a reward * context interaction (see Supplemental Table S6), reflecting Also reaction times were slower, when previously rewarded in the gambling context when contrasted to the neutral context indicating an increased effect of reward on S1 response caution in the gambling context.

**Supplemental Table S6.** Hierarchical general linear model results of S1 RTs with reward and context as fixed effects and subject as random effect.

| S1 RT Model |  |  |  |
| --- | --- | --- | --- |
|  | Estimate | t-Value | p |
| Reward | -0.025 | -2.431 | 0.015* |
| Context | -0.0005 | -0.044 | 0.97 |
| Reward*Context | 0.029 | 2.050 | 0.04* |

### Model free analysis of Stage 2 RTs

**Supplemental Table S7.** Model agnostic analysis of S2 RTs. HGLM with transition and context as fixed effects and subject as random effect.

| S2 RT Model |  |  |  |
| --- | --- | --- | --- |
|  | Estimate | T-statistic | p |
| Transition | 0.13 | 17.227 | < 2e-16*** |
| Context | 0.0004 | 0.07 | 0.94 |
| Transition*Context | 0.02 | 1.804 | 0.07 |

**Supplemental Figure S3.**
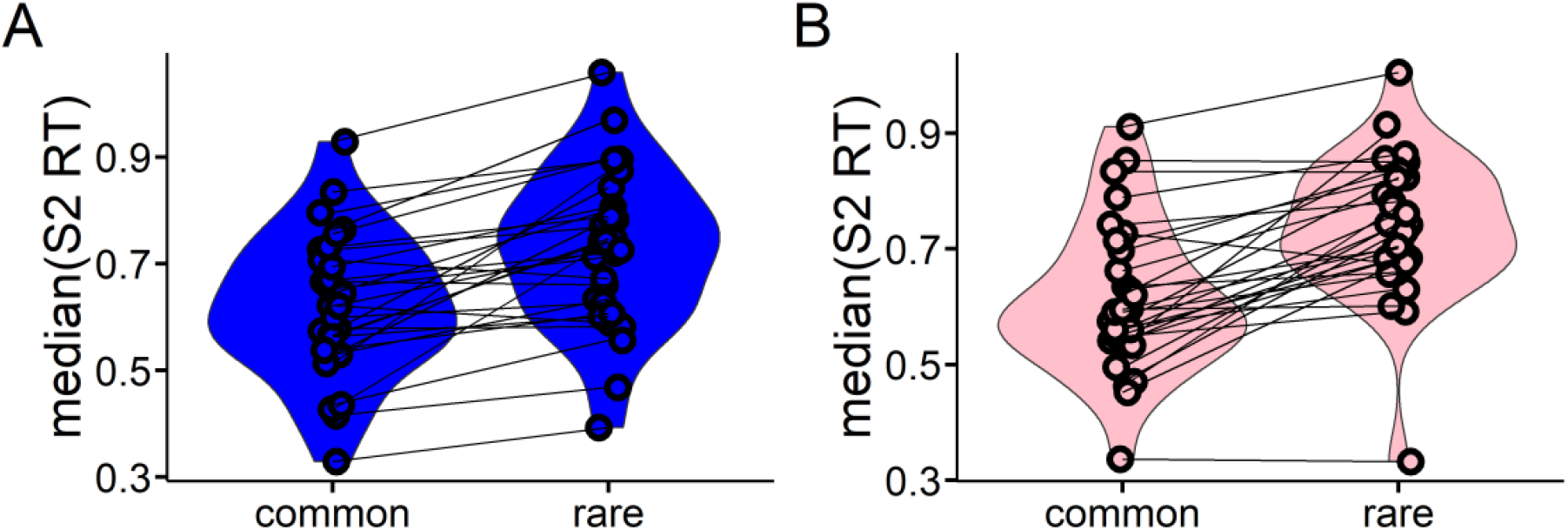
Model free analysis of S2 RTs. RTs were substantially slower following rare transitions, both in the neutral (A) and the gambling context (B), see also Table 4.

### Hybrid model with drift diffusion choice rule

#### Model comparison and validation

Model comparison based on the WAIC (Vehtari et al., 2017) (see Supplemental Table S8) revealed that in the neutral context, a DDM with nonlinear drift-rate scaling DDM_s_ (Fontanesi et al., 2019; Peters & D’Esposito, 2020; Wagner et al., 2020) accounted for the data best when compared to a DDM with linear drift rate scaling (DDM_lin_) (Pedersen et al., 2017) and a null-model without learning (DDM0) (see Supplemental Table S8). The same ranking held for the gambling context.

We next build a full model with group level distributions for the baseline condition (neutral context) and additional s_x_ parameters for each model parameter x, modeling the change in from the neutral to the gambling context. These s_x_ parameters where modeled with Gaussian priors with means of zero (see methods section). The full model reproduced the model ranking (see Supplemental Table S8). We then compared the three DDMs and the softmax model with respect to the proportion of binary choices that they correctly accounted for. As can be seen from see Supplemental Table S9, the DDM_S_ and DDM_lin_ performed numerically on par with the softmax model, whereas the DDM_0_ performed substantially worse. Posterior predictive checks showed that the final model accurately captured the effect of reward differences on second stage RTs and reproduced choice behavior (see Supplemental Figure S4 and S5 below).

**Supplemental Table S8.** Reinforcement learning DDM model comparison using the Widely-Applicable Information Criterion (WAIC) revealed the same model ranking for each condition (neutral or gambling context) as well as for the full model. Scores are WAIC (SE).

|  | Neutral | Gambling | Full model |
| --- | --- | --- | --- |
| **$DDM_0$** | 11808.5 (1549.3) | 10942.1 (1569.2) | 22764.0 (2193.7) |
| **$DDM_{lin}$** | 4616.2 (1897.0) | 4429.0 (1729.9) | 15519.7 (3984.4) |
| **$DDM_s$** | 4357.2 (1935.1) | 4197.2 (1749.7) | 8800.3 (2670.0) |

**Supplemental Table S9.** 2-step task models. Proportions of correctly predicted binary choices (mean [range]) for all models.

|  | Neutral |  | Gambling |  |
| --- | --- | --- | --- | --- |
|  | Stage 1 | Stage 2 | Stage 1 | Stage 2 |
| **DDM<sub>0</sub>** | 0.63 [0.49-1.00] | 0.62 [0.50-0.78] | 0.56 [0.46-0.99] | 0.63 [0.51-0.79] |
| **DDM<sub>lin</sub>** | 0.74 [0.51-1.00] | 0.80 [0.53-0.96] | 0.74 [0.50-0.99] | 0.81 [0.59-0.95] |
| **DDM<sub>s</sub>** | 0.74 [0.49-1.00] | 0.80 [0.55-0.96] | 0.72 [0.47-0.99] | 0.81 [0.59-0.95] |
| **Softmax** | 0.72 [0.42-1.00] | 0.79 [0.49-0.96] | 0.72 [0.45-0.99] | 0.81 [0.56-0.96] |

#### Posterior predictive checks

We conducted posterior predictive checks to evaluate if our different hierarchical models capture both the relationship of RTs and reward differences and the relationship of reward differences and optimal choices. An optimal choice here is defined as a choice for the random walk with highest payout. To this end, we binned all trials into four bins, according to the absolute max(reward) differences in stage 2. For each reward difference bin we then plot the mean observed RTs, as well as the mean simulated RTs across 1000 datasets simulated using our mean parameter estimates for the posterior distributions of the DDM_0_, DDM_lin_, and DDM_S_. We further show the mean observed optimal choices (max[reward]) vs. the mean simulated optimal choices given our mean parameter estimates for the posterior distribution of each model. These results are shown in Supplemental Figures S4 and S5. As can be seen, the DDM_s_ provided the best account of how RTs vary as a function of reward differences. This model outperformed the other models in capturing the relationship of reward differences and optimal choices (Supplemental Figure S5).

**Supplemental Figure S4.**
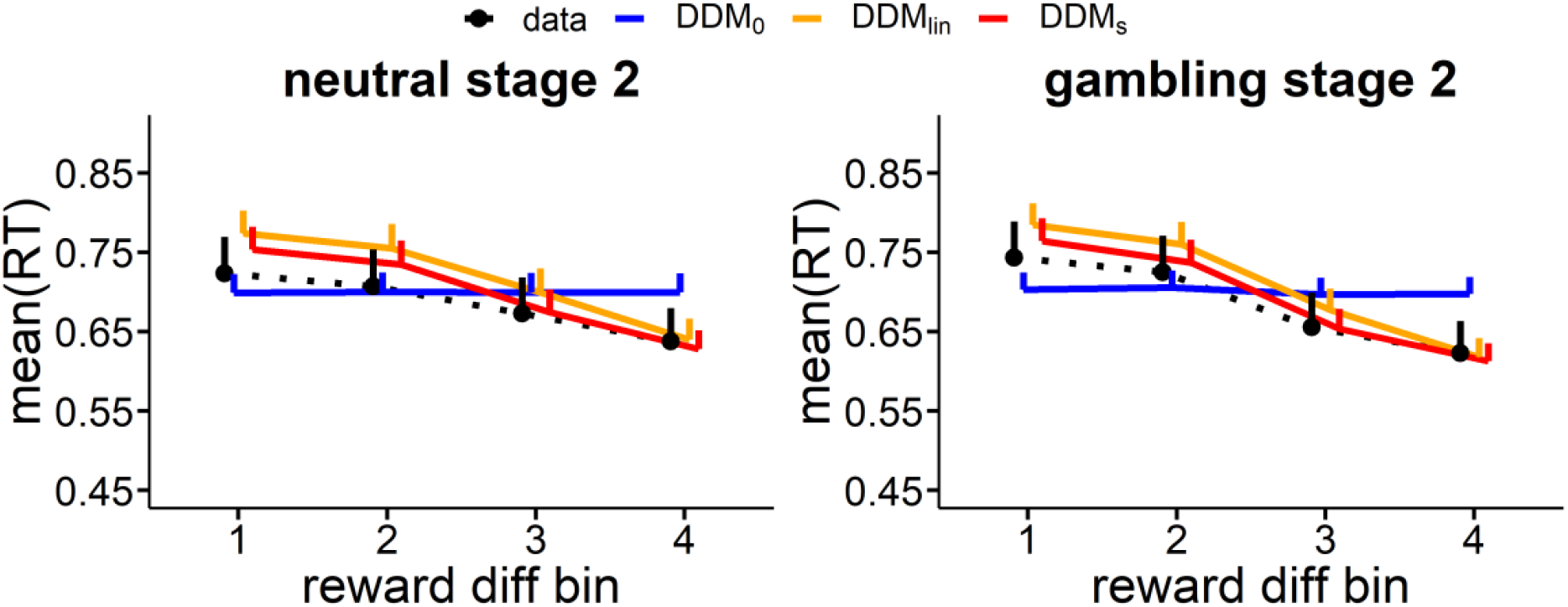
Group level posterior predictive checks. Trials were binned into four equal sized bins according to the absolute difference in *reward* values given S2 reward walks. Plotted are mean observed RTs per bin (data; dashed line) as well model-generated RTs (blue represents DDM_0_; orange represents DDM_lin_; red represents DDM_S_) averaged 1000 simulated datasets simulated from the mean parameter estimates the posterior distribution of each hierarchical model.

**Supplemental Figure S5.**
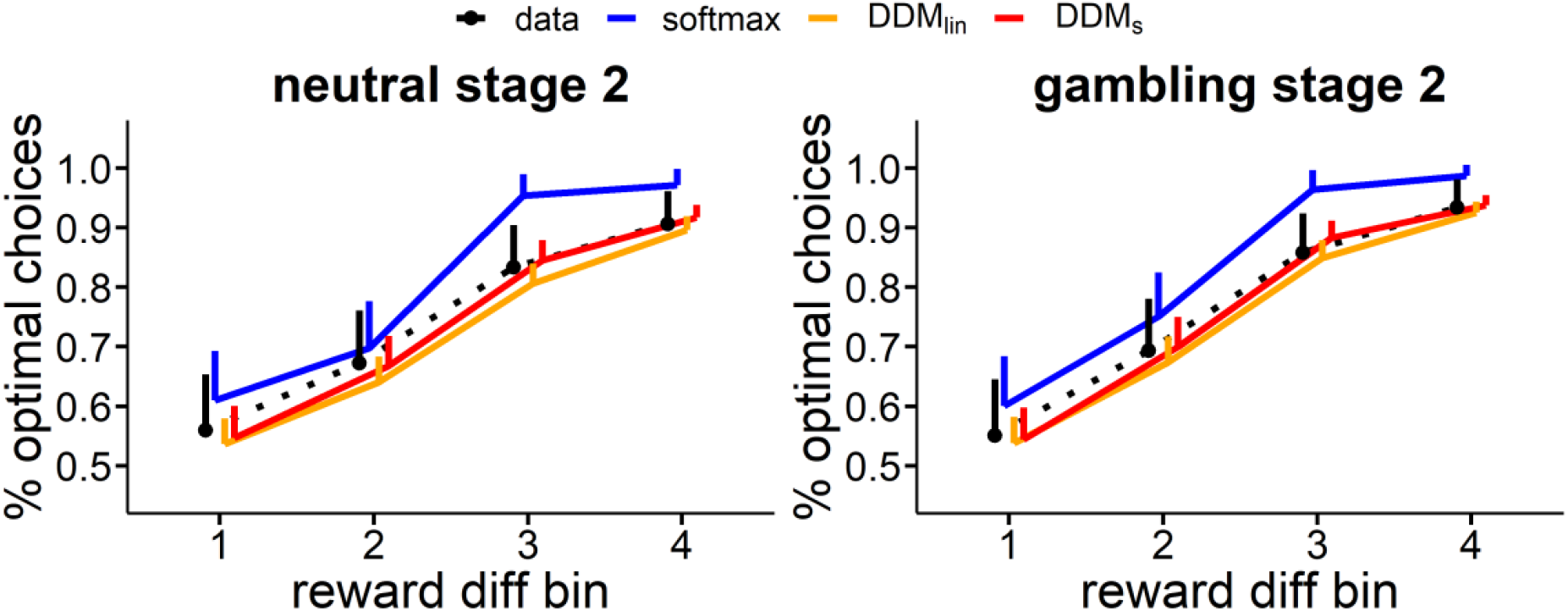
Group level posterior predictive checks. Trials were binned into four equal sized bins according to the absolute difference in *reward* values given S2 reward walks. Plotted are mean optimal choices in % of all choices per bin (data; dashed line) as well model-generated RTs (blue represents DDM_0_; orange represents DDM_lin_; red represents DDM_S_) averaged 1000 simulated datasets simulated from the mean parameter estimates for the posterior distribution of each hierarchical model.

## Supplemental Results 1: Working memory, subjective craving and temporal discounting

### Preregistered analysis

We hypothesized a positive relationship of decision noise parameter and working memory z-score, which we confirmed in each context (neutral condition: r = 0.45, p = 0.013); gambling condition: r = 0.45 p = p = 0.013).

### Exploratory analysis

There was no significant correlation between discount-rate and working memory (r = -0.03, p = 0.89). A further exploratory analysis revealed an association of working memory and drift-rate coefficient. Here, higher working memory capacity was associated with higher drift rate coefficients (neutral condition: r = 0.42, p = 0.02; gambling condition: r = 0.37, p =0.048). The difference in subjective craving from the neutral to the gambling environment, was not significantly associated with the change in the discount-rate (pre-task craving rating: r = 0.28, p = 0.18; post-task craving rating: 0.09, p = 0.68).

## Supplemental Results 2: Working memory, subjective craving and 2-step task performance

### Preregistered analysis

WM z-score was positively but non-significant correlated with MB RL (neutral context: r = 0.27, p = 0.16, gambling context: r = 0.31, p = 0.10). The change in MB RL from neutral to gambling context was not significantly correlated with WM (r = 0.27, p = 0.16).

### Exploratory analysis

We further explored the association of WM capacity and S2 learning rates. This analysis revealed that overall WM capacity was positively associated with baseline (neutral context) (r = 0.57, p = 0.001) and gambling context (r = 0.37, p = 0.048) S2 learning rates. The change in subjective craving from the neutral to the gambling environment, was not significantly associated with the change in MB drift-rate weight (pre-task craving rating: -0.01, p = 0.98; post-task craving rating: r = 0.33, p = 0.14).

**Supplemental Figure S6.**
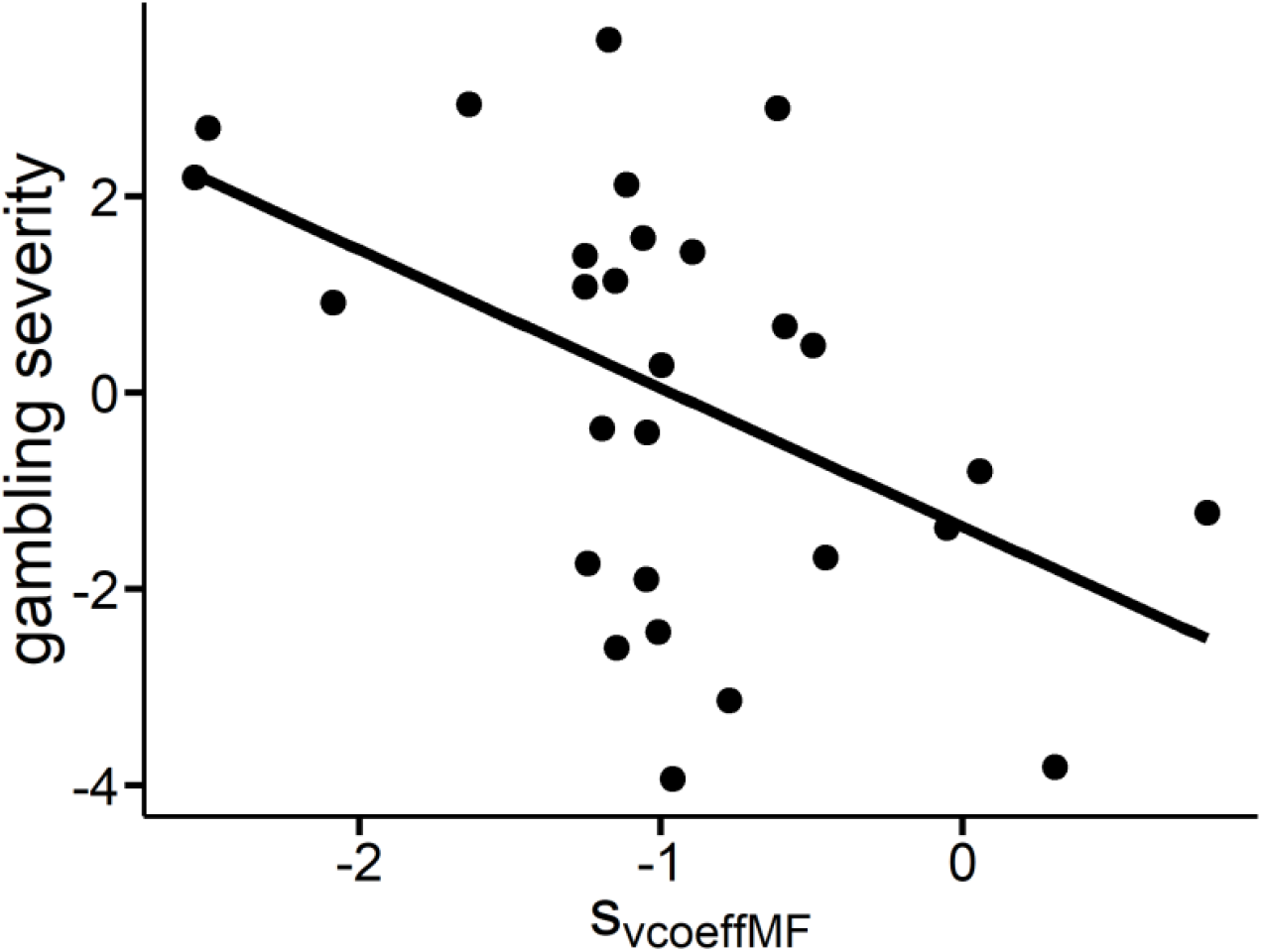
Gambling severity (y-axis; average z-score across DSM, KFG and SOGS) was associated with a greater gambling context related decrease in MF drift-rate weights (*S*_*vcoeffMF*_, r = -0.48, p = 0.009).

## References

American Psychiatric Association. (2013). Diagnostic and statistical manual of mental disorders (5th ed.). Author.

Amlung, M., Marsden, E., Holshausen, K., Morris, V., Patel, H., Vedelago, L., Naish, K. R., Reed, D. D., & McCabe, R. E. (2019). Delay Discounting as a Transdiagnostic Process in Psychiatric Disorders: A Meta-analysis. JAMA Psychiatry, 76(11), 1176–1186. https://doi.org/10.1001/jamapsychiatry.2019.2102

Anderson, G., & Brown, R. I. (1984). Real and laboratory gambling, sensation-seeking and arousal. British Journal of Psychology (London, England : 1953), 75 (Pt 3), 401–410. https://doi.org/10.1111/j.2044-8295.1984.tb01910.x

Anselme, P., & Robinson, M. J. F [Mike J. F.] (2013). What motivates gambling behavior? Insight into dopamine’s role. Frontiers in Behavioral Neuroscience, 7, 182. https://doi.org/10.3389/fnbeh.2013.00182

Antons, S., Brand, M., & Potenza, M. N. (2020). Neurobiology of cue-reactivity, craving, and inhibitory control in non-substance addictive behaviors. Journal of the Neurological Sciences, 415, 116952. https://doi.org/10.1016/j.jns.2020.116952

Association, A. P. (2013). Diagnostic and Statistical Manual of Mental Disorders. American Psychiatric Association. https://doi.org/10.1176/appi.books.9780890425596

Balleine, B. W., & O’Doherty, J. P. (2010). Human and rodent homologies in action control: Corticostriatal determinants of goal-directed and habitual action. Neuropsychopharmacology : Official Publication of the American College of Neuropsychopharmacology, 35(1), 48–69. https://doi.org/10.1038/npp.2009.131

Balodis, I. M., & Potenza, M. N. (2020). Common neurobiological and psychological underpinnings of gambling and substance-use disorders. Progress in Neuro-Psychopharmacology & Biological Psychiatry, 99, 109847. https://doi.org/10.1016/j.pnpbp.2019.109847

Barry J Everitt, & Trevor W Robbins (2005). Neural systems of reinforcement for drug addiction: From actions to habits to compulsion. Nature Neuroscience, 8(11), 1481–1489. https://doi.org/10.1038/nn1579

Beck, A., Steer, R., & Brown, G. (1996). Manual for the Beck Depression Inventory-II (BDI-II).

Berke, J. D. (2018). What does dopamine mean? Nature Neuroscience, 21(6), 787–793. https://doi.org/10.1038/s41593-018-0152-y

Berridge, K. C [K. C.]. (2016). Incentive Motivation and Incentive Salience☆. In J. Stein (Ed.), Reference module in neuroscience and biobehavioral psychology. Elsevier. https://doi.org/10.1016/B978-0-12-809324-5.00342-4

Bickel, W. K., Athamneh, L. N., Basso, J. C., Mellis, A. M., DeHart, W. B., Craft, W. H., & Pope, D. (2019). Excessive discounting of delayed reinforcers as a trans-disease process: Update on the state of the science. Current Opinion in Psychology, 30, 59–64. https://doi.org/10.1016/j.copsyc.2019.01.005

Bickel, W. K., Jarmolowicz, D. P., Mueller, E. T., Koffarnus, M. N., & Gatchalian, K. M. (2012). Excessive discounting of delayed reinforcers as a trans-disease process contributing to addiction and other disease-related vulnerabilities: Emerging evidence. Pharmacology & Therapeutics, 134(3), 287–297. https://doi.org/10.1016/j.pharmthera.2012.02.004

Bickel, W. K., Yi, R., Landes, R. D., Hill, P. F., & Baxter, C. (2011). Remember the future: Working memory training decreases delay discounting among stimulant addicts. Biological Psychiatry, 69(3), 260–265. https://doi.org/10.1016/j.biopsych.2010.08.017

Blaszczynski, A., & Nower, L. (2002). A pathways model of problem and pathological gambling. Addiction (Abingdon, England), 97(5), 487–499. https://doi.org/10.1046/j.1360-0443.2002.00015.x

Clark, L., Boileau, I., & Zack, M. (2019). Neuroimaging of reward mechanisms in Gambling disorder: An integrative review. Molecular Psychiatry, 24(5), 674–693. https://doi.org/10.1038/s41380-018-0230-2

Courtney, K. E., Schacht, J. P., Hutchison, K., Roche, D. J. O., & Ray, L. A. (2016). Neural substrates of cue reactivity: Association with treatment outcomes and relapse. Addiction Biology, 21(1), 3–22. https://doi.org/10.1111/adb.12314

Culbreth, A. J., Westbrook, A [Andrew], Daw, N. D., Botvinick, M., & Barch, D. M. (2016). Reduced model-based decision-making in schizophrenia. Journal of Abnormal Psychology, 125(6), 777–787. https://doi.org/10.1037/abn0000164

da Silva, C. F., & Hare, T. A. (2019). Humans are primarily model-based and not model-free learners in the two-stage task (Vol. 20). https://doi.org/10.1101/682922

Dale, G., Rock, A. J., & Clark, G. I. (2019). Cue-Reactive Imagery Mediates the Relationships of Reward Responsiveness with Both Cue-Reactive Urge to Gamble and Positive Affect in Poker-Machine Gamblers. Journal of Gambling Studies. Advance online publication. https://doi.org/10.1007/s10899-019-09864-x

D’Amour-Horvat, V., & Leyton, M. (2014). Impulsive actions and choices in laboratory animals and humans: Effects of high vs. Low dopamine states produced by systemic treatments given to neurologically intact subjects. Frontiers in Behavioral Neuroscience, 8, 432. https://doi.org/10.3389/fnbeh.2014.00432

Daw, N., Gershman, S., Seymour, B., Dayan, P [Peter], & Dolan, R. J. (2011). Model-Based Influences on Humans’ Choices and Striatal Prediction Errors. Neuron, 69(6), 1204–1215. https://doi.org/10.1016/j.neuron.2011.02.027

Dixon, M [Mark], Ghezzi, P., Lyons, C., & Wilson, G. (2006). Gambling: Behavior Theory, Research, and Application. New Harbinger Publications.

Dixon, M [Mark.], Jacobs, E. A., & Sanders, S. (2006). CONTEXTUAL CONTROL OF DELAY DISCOUNTING BY PATHOLOGICAL GAMBLERS. Journal of Applied Behavior Analysis, 39(4), 413–422. https://doi.org/10.1901/jaba.2006.173-05

Dixon, M [Mark.], Marley, J., & Jacobs, E. A. (2003). Delay discounting by pathological gamblers. Journal of Applied Behavior Analysis, 36(4), 449–458. https://doi.org/10.1901/jaba.2003.36-449

Doll, B. B., Simon, D. A., & Daw, N. D. (2012). The ubiquity of model-based reinforcement learning. Current Opinion in Neurobiology, 22(6), 1075–1081. https://doi.org/10.1016/j.conb.2012.08.003

Falkai, P. (2015). Diagnostisches und statistisches Manual psychischer Störungen: DSM-5. Hogrefe.

Faul, F., Erdfelder, E., Lang, A.-G., & Buchner, A. (2007). G*power 3: A flexible statistical power analysis program for the social, behavioral, and biomedical sciences. Behavior Research Methods, 39(2), 175–191. https://doi.org/10.3758/BF03193146

Fauth-Bühler, M., Mann, K., & Potenza, M. N. (2017). Pathological gambling: A review of the neurobiological evidence relevant for its classification as an addictive disorder. Addiction Biology, 22(4), 885–897. https://doi.org/10.1111/adb.12378

FitzGerald, T. H. B., Dolan, R. J., & Friston, K. (2015). Dopamine, reward learning, and active inference. Frontiers in Computational Neuroscience, 9, 136. https://doi.org/10.3389/fncom.2015.00136

Flagel, S. B., Akil, H., & Robinson, T. E [Terry E.] (2009). Individual differences in the attribution of incentive salience to reward-related cues: Implications for addiction. Neuropharmacology, 56 Suppl 1, 139–148. https://doi.org/10.1016/j.neuropharm.2008.06.027

Fontanesi, L., Gluth, S., Spektor, M. S., & Rieskamp, J. (2019). A reinforcement learning diffusion decision model for value-based decisions. Psychonomic Bulletin & Review, 26(4), 1099–1121. https://doi.org/10.3758/s13423-018-1554-2

Forstmann, B. U., Ratcliff, R., & Wagenmakers, E.-J. (2016). Sequential Sampling Models in Cognitive Neuroscience: Advantages, Applications, and Extensions. Annual Review of Psychology, 67, 641–666. https://doi.org/10.1146/annurev-psych-122414-033645

Frank, M. J [Michael J.], & O’Reilly, R. C. (2006). A mechanistic account of striatal dopamine function in human cognition: Psychopharmacological studies with cabergoline and haloperidol. Behavioral Neuroscience, 120(3), 497–517. https://doi.org/10.1037/0735-7044.120.3.497

Genauck, A., Andrejevic, M., Brehm, K., Matthis, C., Heinz, A., Weinreich, A., Kathmann, N., & Romanczuk-Seiferth, N. (2020). Cue-induced effects on decision-making distinguish subjects with gambling disorder from healthy controls. Addiction Biology, 25(6), e12841. https://doi.org/10.1111/adb.12841

Gershman, S. J., & Bhui, R. (2020). Rationally inattentive intertemporal choice. Nature Communications, 11(1), 3365. https://doi.org/10.1038/s41467-020-16852-y

Gillan, C. M., Kalanthroff, E., Evans, M., Weingarden, H. M., Jacoby, R. J., Gershkovich, M., Snorrason, I., Campeas, R., Cervoni, C., Crimarco, N. C., Sokol, Y., Garnaat, S. L., McLaughlin, N. C. R., Phelps, E. A., Pinto, A., Boisseau, C. L., Wilhelm, S., Daw, N. D., & Simpson, H. B [H. B.] (2020). Comparison of the Association Between Goal-Directed Planning and Self-reported Compulsivity vs Obsessive-Compulsive Disorder Diagnosis. JAMA Psychiatry, 77(1), 77–85. https://doi.org/10.1001/jamapsychiatry.2019.2998

Gillan, C. M., Kosinski, M., Whelan, R., Phelps, E. A., & Daw, N. D. (2016). Characterizing a psychiatric symptom dimension related to deficits in goal-directed control. ELife, 5. https://doi.org/10.7554/eLife.11305

Hautzinger, M., Keller, F., & Kühner, C. (2009). BDI-II. Beck-Depressions-Inventar. Revision. 2, Auflage.

Heatherton, T. F., Kozlowski, L. T., Frecker, R. C., & Fagerstrom, K. O. (1991). The Fagerström Test for Nicotine Dependence: A revision of the Fagerstrom Tolerance Questionnaire. British Journal of Addiction, 86(9), 1119–1127.

Joukhador, J., Blaszczynski, A., & Maccallum, F. (2004). Superstitious beliefs in gambling among problem and non-problem gamblers: Preliminary data. Journal of Gambling Studies, 20(2), 171–180. https://doi.org/10.1023/B:JOGS.0000022308.27774.2b

Kleiner, M., Brainard, D., & Pelli, D. (2007). What’s new in Psychtoolbox-3? https://pure.mpg.de/rest/items/item_1790332/component/file_3136265/content

Kool, W., Cushman, F. A., & Gershman, S. J. (2016). When Does Model-Based Control Pay Off? PLOS Computational Biology, 12(8), e1005090. https://doi.org/10.1371/journal.pcbi.1005090

Kroemer, N. B., Lee, Y., Pooseh, S., Eppinger, B., Goschke, T., & Smolka, M. N. (2019). L-DOPA reduces model-free control of behavior by attenuating the transfer of value to action. NeuroImage, 186, 113–125. https://doi.org/10.1016/j.neuroimage.2018.10.075

Leeman, R. F., & Potenza, M. N. (2012). Similarities and differences between pathological gambling and substance use disorders: A focus on impulsivity and compulsivity. Psychopharmacology, 219(2), 469–490. https://doi.org/10.1007/s00213-011-2550-7

Lempert, K. M., & Phelps, E. A. (2016). The Malleability of Intertemporal Choice. Trends in Cognitive Sciences, 20(1), 64–74. https://doi.org/10.1016/j.tics.2015.09.005

Lempert, K. M., Steinglass, J. E., Pinto, A., Kable, J. W., & Simpson, H. B [Helen Blair] (2019). Can delay discounting deliver on the promise of RDoC? Psychological Medicine, 49(2), 190–199. https://doi.org/10.1017/S0033291718001770

Lesieur, H. R., & Blume, S. B. (1987). The South Oaks Gambling Screen (SOGS): A new instrument for the identification of pathological gamblers. American Journal of Psychiatry, 144(9).

Lobo, D. S. S., Aleksandrova, L., Knight, J., Casey, D. M., el-Guebaly, N., Nobrega, J. N., & Kennedy, J. L. (2015). Addiction-related genes in gambling disorders: New insights from parallel human and pre-clinical models. Molecular Psychiatry, 20(8), 1002. https://doi.org/10.1038/mp.2014.113

MacKillop, J., Amlung, M. T., Few, L. R., Ray, L. A., Sweet, L. H., & Munafò, M. R. (2011). Delayed reward discounting and addictive behavior: A meta-analysis. Psychopharmacology, 216(3), 305–321. https://doi.org/10.1007/s00213-011-2229-0

Martyn Plummer. (2003). Jags: A Program for Analysis of Bayesian Graphical Models Using Gibbs Sampling. In M. Plummer (Ed.), JAGS: a program for analysis of Bayesian graphical models using Gibbs sampling.: Proceedings of the 3rd International Workshop on Distributed Statistical Computing (DSC 2003). 2003.

Miedl, S. F., Büchel, C., & Peters, J. (2014). Cue-induced craving increases impulsivity via changes in striatal value signals in problem gamblers. The Journal of Neuroscience : The Official Journal of the Society for Neuroscience, 34(13), 4750–4755. https://doi.org/10.1523/JNEUROSCI.5020-13.2014

Miedl, S. F., Peters, J., & Büchel, C. (2012). Altered neural reward representations in pathological gamblers revealed by delay and probability discounting. Archives of General Psychiatry, 69(2), 177–186. https://doi.org/10.1001/archgenpsychiatry.2011.1552

Mikhael, J. G., Lai, L., & Gershman, S. J. (2021). Rational inattention and tonic dopamine. PLOS Computational Biology, 17(3), e1008659. https://doi.org/10.1371/journal.pcbi.1008659

Moeller, S. J., & Paulus, M. P. (2018). Toward biomarkers of the addicted human brain: Using neuroimaging to predict relapse and sustained abstinence in substance use disorder. Progress in Neuro-Psychopharmacology & Biological Psychiatry, 80(Pt B), 143–154. https://doi.org/10.1016/j.pnpbp.2017.03.003

Muggleton, N., Parpart, P., Newall, P., Leake, D., Gathergood, J., & Stewart, N. (2021). The association between gambling and financial, social and health outcomes in big financial data. Nature Human Behaviour. Advance online publication. https://doi.org/10.1038/s41562-020-01045-w

Myerson, J., Green, L., & Warusawitharana, M. (2001). Area under the curve as a measure of discounting. Journal of the Experimental Analysis of Behavior, 76(2), 235–243. https://doi.org/10.1901/jeab.2001.76-235

Nelson, L. D., Strickland, C., Krueger, R. F., Arbisi, P. A., & Patrick, C. J. (2016). Neurobehavioral Traits as Transdiagnostic Predictors of Clinical Problems. Assessment, 23(1), 75–85. https://doi.org/10.1177/1073191115570110

Ott, T., & Nieder, A. (2019). Dopamine and Cognitive Control in Prefrontal Cortex. Trends in Cognitive Sciences, 23(3), 213–234. https://doi.org/10.1016/j.tics.2018.12.006

Otto, A. R., Skatova, A., Madlon-Kay, S., & Daw, N. D. (2015). Cognitive control predicts use of model-based reinforcement learning. Journal of Cognitive Neuroscience, 27(2), 319–333. https://doi.org/10.1162/jocn_a_00709

Patzelt, E. H., Kool, W., Millner, A. J., & Gershman, S. J. (2019). Incentives Boost Model-Based Control Across a Range of Severity on Several Psychiatric Constructs. Biological Psychiatry, 85(5), 425–433. https://doi.org/10.1016/j.biopsych.2018.06.018

Pedersen, M. L., Frank, M. J [Michael J.], & Biele, G. (2017). The drift diffusion model as the choice rule in reinforcement learning. Psychonomic Bulletin & Review, 24(4), 1234–1251. https://doi.org/10.3758/s13423-016-1199-y

Peters, J., & Büchel, C. (2010). Episodic future thinking reduces reward delay discounting through an enhancement of prefrontal-mediotemporal interactions. Neuron, 66(1), 138–148. https://doi.org/10.1016/j.neuron.2010.03.026

Peters, J., & Büchel, C. (2011). The neural mechanisms of inter-temporal decision-making: Understanding variability. Trends in Cognitive Sciences, 15(5), 227–239. https://doi.org/10.1016/j.tics.2011.03.002

Peters, J., & D’Esposito, M. (2020). The drift diffusion model as the choice rule in intertemporal and risky choice: A case study in medial orbitofrontal cortex lesion patients and controls. PLOS Computational Biology, 16(4), e1007615. https://doi.org/10.1371/journal.pcbi.1007615

Petry, J., & Baulig, T. (1996). KFG: Kurzfragebogen zum Glücksspielverhalten. Psychotherapie Der Glücksspielsucht, 300–302.

Petry, N. M. (2010). Pathological gambling and the DSM-V. International Gambling Studies, 10(2), 113–115. https://doi.org/10.1080/14459795.2010.501086

Petzold, J., Kienast, A., Lee, Y., Pooseh, S., London, E. D., Goschke, T., & Smolka, M. N. (2019). Baseline impulsivity may moderate L-DOPA effects on value-based decisionmaking. Scientific Reports, 9(1), 5652. https://doi.org/10.1038/s41598-019-42124-x

Pine, A., Shiner, T., Seymour, B., & Dolan, R. J. (2010). Dopamine, time, and impulsivity in humans. The Journal of Neuroscience : The Official Journal of the Society for Neuroscience, 30(26), 8888–8896. https://doi.org/10.1523/JNEUROSCI.6028-09.2010

Raylu, N., & Oei, T. P. S. (2004). The Gambling Related Cognitions Scale (GRCS): Development, confirmatory factor validation and psychometric properties. Addiction (Abingdon, England), 99(6), 757–769.

Redick, T. S., Broadway, J. M., Meier, M. E., Kuriakose, P. S., Unsworth, N., Kane, M. J., & Engle, R. W. (2012). Measuring Working Memory Capacity With Automated Complex Span Tasks. European Journal of Psychological Assessment, 28(3), 164–171. https://doi.org/10.1027/1015-5759/a000123

Reynolds, B. (2006). A review of delay-discounting research with humans: Relations to drug use and gambling. Behavioural Pharmacology, 17(8), 651–667. https://doi.org/10.1097/FBP.0b013e3280115f99

Robbins, T. W., & Everitt, B. J. (1999). Drug addiction: Bad habits add up. Nature, 398(6728), 567–570. https://doi.org/10.1038/19208

Robinson, M. J. F [M. J. F.], Fischer, A. M., Ahuja, A., Lesser, E. N., & Maniates, H. (2016). Roles of “Wanting” and “Liking” in Motivating Behavior: Gambling, Food, and Drug Addictions. Current Topics in Behavioral Neurosciences, 27, 105–136. https://doi.org/10.1007/7854_2015_387

Robinson, T., & Berridge, K. C [Kent C.] (1993). The neural basis of drug craving: An incentive-sensitization theory of addiction. Brain Research Reviews, 18(3), 247–291. https://doi.org/10.1016/0165-0173(93)90013-P

Robinson, T. E., & Berridge, K. C [K. C.] (2001). Incentive-sensitization and addiction. Addiction (Abingdon, England), 96(1), 103–114. https://doi.org/10.1046/j.1360-0443.2001.9611038.x

Robinson, T. E [Terry E.], & Berridge, K. C [Kent C.] (2008). Review. The incentive sensitization theory of addiction: Some current issues. Philosophical Transactions of the Royal Society B: Biological Sciences, 363(1507), 3137–3146. https://doi.org/10.1098/rstb.2008.0093

Rösch, S. A., Stramaccia, D. F., & Benoit, R. G. (2021). Promoting farsighted decisions via episodic future thinking: A meta-analysis. https://doi.org/10.31234/osf.io/53ju2

Saunders, J. B., Aasland, O. G., Babor, T. F., La Fuente, J. R. de, & Grant, M. (1993). Development of the alcohol use disorders identification test (AUDIT): WHO collaborative project on early detection of persons with harmful alcohol consumption-II. Addiction (Abingdon, England), 88(6), 791–804.

Sebold, M., Deserno, L., Nebe, S [Stephan], Nebe, S [Stefan], Schad, D. J., Garbusow, M., Hägele, C., Keller, J., Jünger, E., Kathmann, N., Smolka, M. N., Smolka, M., Rapp, M. A., Schlagenhauf, F., Heinz, A., & Huys, Q. J. M. (2014). Model-based and model-free decisions in alcohol dependence. Neuropsychobiology, 70(2), 122–131. https://doi.org/10.1159/000362840

Shahar, N., Hauser, T. U., Moutoussis, M., Moran, R., Keramati, M., & Dolan, R. J. (2019). Improving the reliability of model-based decision-making estimates in the two-stage decision task with reaction-times and drift-diffusion modeling. PLoS Computational Biology, 15(2), e1006803. https://doi.org/10.1371/journal.pcbi.1006803

Sharp, M. E., Foerde, K., Daw, N. D., & Shohamy, D. (2016). Dopamine selectively remediates ‘model-based’ reward learning: A computational approach. Brain : A Journal of Neurology, 139(Pt 2), 355–364. https://doi.org/10.1093/brain/awv347

Shiner, T., Seymour, B., Wunderlich, K., Hill, C., Bhatia, K. P., Dayan, P [Peter], & Dolan, R. J. (2012). Dopamine and performance in a reinforcement learning task: Evidence from Parkinson’s disease. Brain : A Journal of Neurology, 135(Pt 6), 1871–1883. https://doi.org/10.1093/brain/aws083

Singer, B. F., Anselme, P., Robinson, M. J. F [Mike J. F.], & Vezina, P. (2020). An overview of commonalities in the mechanisms underlying gambling and substance use disorders. Progress in Neuro-Psychopharmacology & Biological Psychiatry, 101, 109944. https://doi.org/10.1016/j.pnpbp.2020.109944

Stan Development Team. (2020). Stan (Version 2.26) [Computer software]. https://mc-stan.org

Toyama, A., Katahira, K., & Ohira, H. (2017). A simple computational algorithm of model-based choice preference. Cognitive, Affective & Behavioral Neuroscience, 17(4), 764–783. https://doi.org/10.3758/s13415-017-0511-2

Toyama, A., Katahira, K., & Ohira, H. (2019). Reinforcement Learning With Parsimonious Computation and a Forgetting Process. Frontiers in Human Neuroscience, 13, 153. https://doi.org/10.3389/fnhum.2019.00153

van den Noort, M., Bosch, P., Haverkort, M., & Hugdahl, K. (2008, January 21). A Standard Computerized Version of the Reading Span Test in Different Languages. Hogrefe & Huber Publishers. https://econtent.hogrefe.com/doi/abs/10.1027/1015-5759.24.1.35

van Gaalen, M. M., van Koten, R., Schoffelmeer, A. N. M., & Vanderschuren, L. J. M. J. (2006). Critical involvement of dopaminergic neurotransmission in impulsive decision making. Biological Psychiatry, 60(1), 66–73. https://doi.org/10.1016/j.biopsych.2005.06.005

Vehtari, A., Gelman, A., & Gabry, J. (2017). Practical Bayesian model evaluation using leave-one-out cross-validation and WAIC. Statistics and Computing, 27(5), 1413–1432. https://doi.org/10.1007/s11222-016-9696-4

Voon, V [V.], Derbyshire, K., Rück, C., Irvine, M. A., Worbe, Y., Enander, J., Schreiber, L. R. N., Gillan, C., Fineberg, N. A., Sahakian, B. J., Robbins, T. W., Harrison, N. A., Wood, J., Daw, N. D [N. D.], Dayan, P [P.], Grant, J. E., & Bullmore, E. T. (2015a). Disorders of compulsivity: A common bias towards learning habits. Molecular Psychiatry, 20(3), 345–352. https://doi.org/10.1038/mp.2014.44

Voon, V [V.], Derbyshire, K., Rück, C., Irvine, M. A., Worbe, Y., Enander, J., Schreiber, L. R. N., Gillan, C., Fineberg, N. A., Sahakian, B. J., Robbins, T. W., Harrison, N. A., Wood, J., Daw, N. D [N. D.], Dayan, P [P.], Grant, J. E., & Bullmore, E. T. (2015b). Disorders of compulsivity: A common bias towards learning habits. Molecular Psychiatry, 20(3), 345–352. https://doi.org/10.1038/mp.2014.44

Voon, V [Valerie], Reiter, A., Sebold, M., & Groman, S. (2017). Model-Based Control in Dimensional Psychiatry. Biological Psychiatry, 82(6), 391–400. https://doi.org/10.1016/j.biopsych.2017.04.006

Wagner, B., Clos, M., Sommer, T., & Peters, J. (2020). Dopaminergic Modulation of Human Intertemporal Choice: A Diffusion Model Analysis Using the D2-Receptor Antagonist Haloperidol. The Journal of Neuroscience : The Official Journal of the Society for Neuroscience, 40(41), 7936–7948. https://doi.org/10.1523/JNEUROSCI.0592-20.2020

Westbrook, A [A.], Frank, M [MJ.], & Cools, R. (2021). A mosaic of cost-benefit control over cortico-striatal circuitry. Trends in Cognitive Sciences. Advance online publication. https://doi.org/10.1016/j.tics.2021.04.007

Westbrook, A [A.], van den Bosch, R., Määttä, J. I., Hofmans, L., Papadopetraki, D., Cools, R., & Frank, M. J [M. J.] (2020). Dopamine promotes cognitive effort by biasing the benefits versus costs of cognitive work. Science, 367(6484), 1362–1366. https://doi.org/10.1126/science.aaz5891

Wiehler, A., & Peters, J. (2015). Reward-based decision making in pathological gambling: The roles of risk and delay. Neuroscience Research, 90, 3–14. https://doi.org/10.1016/j.neures.2014.09.008

Wilke, A., Scheibehenne, B., Gaissmaier, W., McCanney, P., & Barrett, H. C. (2014). Illusionary pattern detection in habitual gamblers. Evolution and Human Behavior, 35(4), 291–297. https://doi.org/10.1016/j.evolhumbehav.2014.02.010

Wunderlich, K., Smittenaar, P., & Dolan, R. J. (2012). Dopamine enhances model-based over model-free choice behavior. Neuron, 75(3), 418–424. https://doi.org/10.1016/j.neuron.2012.03.042

Wyckmans, F., Otto, A. R., Sebold, M., Daw, N., Bechara, A., Saeremans, M., Kornreich, C., Chatard, A., Jaafari, N., & Noël, X. (2019). Reduced model-based decision-making in gambling disorder. Scientific Reports, 9(1), 19625. https://doi.org/10.1038/s41598-019-56161-z

